# Flow zoometry of *Drosophila*

**DOI:** 10.1101/2024.04.04.588032

**Authors:** Walker Peterson, Joshua Arenson, Soichiro Hata, Laura Kacenauskaite, Tsubasa Kobayashi, Takuya Otsuka, Hanqing Wang, Yayoi Wada, Kotaro Hiramatsu, Zhikai He, Jean-Emmanuel Clement, Chenqi Zhang, Chenglang Hu, Phillip McCann, Hayato Kanazawa, Yuzuki Nagasaka, Hiroyuki Uechi, Yuh Watanabe, Ryodai Yamamura, Mika Hayashi, Yuta Nakagawa, Kangrui Huang, Hiroshi Kanno, Yuqi Zhou, Tianben Ding, Maik Herbig, Shimpei Makino, Shunta Nonaga, Ryosuke Takami, Oguz Kanca, Koji Tabata, Satoshi Amaya, Kotaro Furusawa, Kenichi Ishii, Kazuo Emoto, Fumihito Arai, Ross Cagan, Dino Di Carlo, Tatsushi Igaki, Erina Kuranaga, Shinya Yamamoto, Hugo J Bellen, Tamiki Komatsuzaki, Masahiro Sonoshita, Keisuke Goda

## Abstract

*Drosophila* serves as a highly valuable model organism across numerous fields including genetics, immunology, neuroscience, cancer biology, and developmental biology. Central to *Drosophila*-based biological research is the ability to perform comprehensive genetic or chemical screens. However, this research is often limited by its dependence on laborious manual handling and analysis, making it prone to human error and difficult to discern statistically significant or rare events amid the noise of individual variations resulting from genetic and environmental factors. In this article we present flow zoometry, a whole-animal equivalent of flow cytometry for large-scale, individual-level, high-content screening of *Drosophila*. Our flow zoometer automatically clears the tissues of *Drosophila melanogaster*, captures three-dimensional (3D) multi-color fluorescence tomograms of single flies with single-cell volumetric resolution at an unprecedented throughput of over 1,000 animals within 48 hours (24 hr for clearing; 24 hr for imaging), and performs AI-enhanced data-driven analysis – a task that would traditionally take months or years with manual techniques. To demonstrate its broad applications, we employed the flow zoometer in various laborious screening assays, including those in toxicology, genotyping, and tumor screening. Flow zoometry represents a pivotal evolution in high-throughput screening technology: previously from molecules to cells, now from cells to whole animals. This advancement serves as a foundational platform for “statistical spatial biology”, to improve empirical precision and enable serendipitous discoveries across various fields of biology.

## INTRODUCTION

*Drosophila*, a highly valuable model organism, is used extensively in diverse biological fields such as genetics, immunology, neuroscience, cancer biology, and developmental biology^1–10^. By virtue of significant gene homology, genetic findings from *Drosophila* frequently translate well to mammals, including humans. Major advantages of *Drosophila* compared to other model organisms include its high generational turnover, cost-effectiveness, and ability to perform comprehensive whole-animal genetic or chemical screens^11–21^. Furthermore, the ease of *Drosophila* genetic engineering facilitates the elucidation of specific gene functions at a systemic level, thereby enhancing its versatility in biological research^22^. Lastly, *Drosophila* research avoids many of the ethical and regulatory complications inherent to mammalian testing^23^. These collective advantages render *Drosophila* an attractive model organism for researchers across diverse fields.

However, the full potential of *Drosophila* has yet to be realized. Comprehensive screens are laborious, which makes it difficult to implement statistically significant sample sizes required to detect subtle or infrequent biological phenomena amidst the noise of individual-level variations resulting from genetic and environmental differences^24–26^. For instance, the common practice of manual dissection of *Drosophila* for intricate examination of internal tissues is time-consuming, labor-intensive, and reliant on operator skill, posing scalability issues that curtail accuracy and exploration in biological research. While cell-based research has seen advancements with high-throughput technologies^27,28^, there is a dearth of similar innovations in the domain of whole animals that specifically contribute to multi-organ network analysis. While several methods for large-scale screening of *Drosophila* have been proposed^29–31^, these mainly focus on embryonic stages and are constrained to the analysis of embryogenesis; since embryos do not have fully developed tissues and are incapable of consuming chemical-containing food, embryonic-stage methods cannot be used to explore complex anatomical systems or study diseases and therapeutics. Although several techniques have been developed to obtain whole-body images of *Drosophila* larvae^32,33^, these are not compatible with large-scale imaging at a high throughput.

Flow zoometry is a foundational technology that addresses multiple key challenges and enables precise whole-body characterization of *Drosophila melanogaster* third instar larvae (the final stage of larval development known as L3, with fully formed anatomical systems) on an unprecedented scale. It significantly enhances the utility of *Drosophila* as a model organism, offering a level of comprehensive analysis previously unattainable in whole animals. With our flow zoometer, we can produce 3D multi-color fluorescence tomograms with single-cell volumetric resolution, covering 6 Gigavoxels and 12 GB of data for each animal, processing tissue-cleared larvae at a rate greater than 1,000 animals per 24 hr (after an automated 24-hr tissue-clearing process), corresponding to a throughput of one animal per minute. This efficiency expands the ability to explore biological questions with whole-animal sophistication and significant statistical power.

## RESULTS

### Principles of flow zoometry

The flow zoometer is based on the integration of five key components, detailed in Figure 1 (refer to Supplementary Video 1 for an overview of its functionality, Supplementary Video 2 for the flow zoometer in operation, Supplementary Figure 1 and Supplementary Figure 2 for detailed schematics of the flow zoometer, and Supplementary Figure 3 for pictures of the flow zoometer).

**Figure 1.**
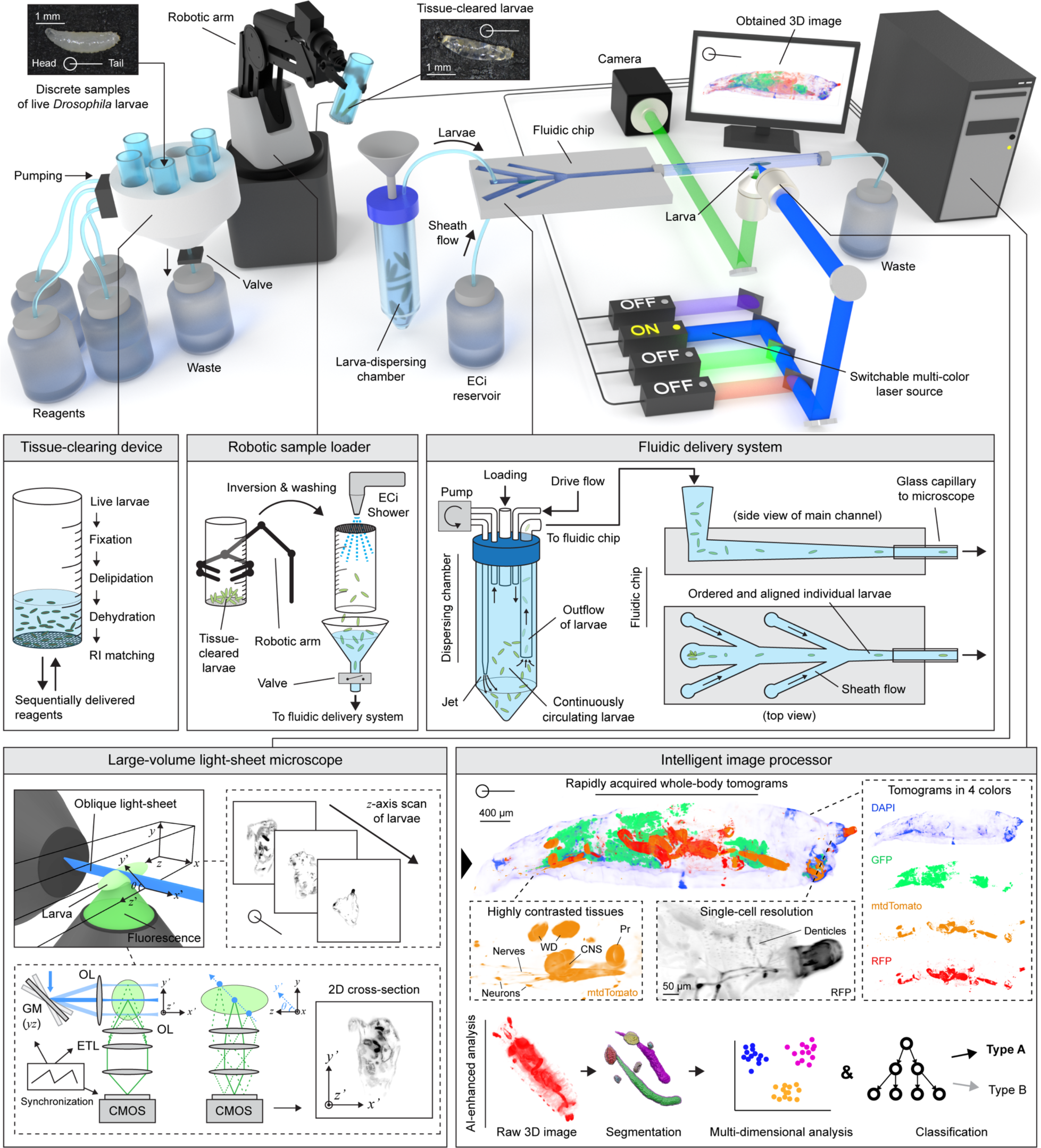
Flow zoometry of *Drosophila*. The fully automated flow zoometer, comprised of five key components, facilitates a streamlined workflow for processing, imaging, and analyzing *Drosophila* larvae (refer to Supplementary Videos 1 and 2 for a comprehensive demonstration of the flow zoometer, Supplementary Figures 1 and 2 for intricate schematics, and Supplementary Figure 3 for pictures of the flow zoometer). The process initiates with the tissue-clearing device, where multiple live *Drosophila* larvae are fixed, delipidated, dehydrated, and refractive index matched. The robotic sample loader then transfers these tissue-cleared larvae to the fluidic delivery system. This system, with its dispersing chamber and fluidic chip, efficiently orders, orients, and delivers each larva to the large-volume LSM via a glass capillary. Subsequently, each larva is immobilized in the LSM, and a stack of 2D fluorescence cross-sections of the larva’s short axis is captured by translating the larva along its long axis through an oblique light-sheet (highlighted in the upper-right inset). The captured tomograms are relayed to the intelligent image processor which, through a series of computational steps, produces multi-colored 3D images of whole larvae with highly contrasted tissues at single-cell resolution. Additionally, it performs data-driven analysis including segmentation, multi-dimensional analysis, and AI-based classification. Remarkably, this fully automated platform can clear, measure, and analyze over 1,000 *Drosophila* L3 larvae within a span of 48 hr (24 hr for clearing; 24 hr for imaging). OL: objective lens; GM: galvanometric mirrors; ETL: electrotunable lens; WD: wing disc; Pr: proventriculus; CNS: central nervous system.

The first component is a fully automated tissue-clearing device capable of fixing and clearing over 1,000 *Drosophila* L3 larvae within 24 hr. Samples of live or fixed larvae in solution are loaded into mesh-bottomed sample vials to filter out liquids and unwanted debris. After loading, reagents are sequentially delivered to the larvae for automated clearing. The device employs an ethyl cinnamate (ECi)-based tissue-clearing protocol^34^ for simplicity and safety. Once the tissue-clearing process is complete, the cleared larvae are suspended in ECi and ready for storage or imaging (see Supplementary Figure 4 for images of tissue-cleared larvae produced by the automated tissue-clearing device).

The second component is a robotic sample loader. Using a robotic arm, sample vials filled with tissue-cleared larvae are transported from the tissue-clearing device to a fluidic delivery system. The vials are inverted over a funnel, then an ECi shower system is used to gently flush the larvae from the vial into the funnel. The larvae then seamlessly flow from the funnel into the third component of the flow zoometer, the fluidic delivery system. To maintain a hermetic seal between the funnel and the fluidic delivery system, a valve is securely closed after the loading process.

The third component is an entirely ECi-based fluidic delivery system. It operates in two phases to isolate and transport individual larvae: the first utilizes a dispersing chamber, and the second a fluidic chip. The dispersing chamber features a jet that ensures continuously circulating larvae in the bottom half of the chamber. Concurrently, a drive flow introduced at the chamber’s top causes outflow of larvae toward the fluidic chip through a pickup tube positioned near the middle of the chamber. The fluidic chip is designed with five sheath flows that converge on the main channel to further separate the larvae. These sheath flows align each larva in accordance with the flow direction, ensuring individual larvae enter a 2-mm square glass capillary for imaging without causing blockages. Within the capillary, an incoming larva is identified by a laser trigger system. When a larva is detected, all fluid movement in the fluidic delivery system is instantly halted by closing valves immediately preceding and following the capillary and deactivating all pumps, effectively immobilizing the larva within the capillary.

The fourth component is a large-volume light-sheet microscope (LSM), tailored to accommodate the substantial size of an entire *Drosophila* L3 larva, with an imaging volume of up to 2 mm × 2 mm × 40 mm. The LSM consists of an excitation arm equipped with a switchable multi-color laser source (up to 5 colors), a switcher for Gaussian or Bessel beams, a 2-axis galvanometric scanning mirror, relay lenses, and an objective lens. The sample is scanned by the galvanometer mirrors, forming a 2D oblique light-sheet at a 27° angle relative to the objective lens of the detection arm. The detection arm includes an objective lens, an electrotunable lens, relay lenses, a 6-color fluorescence filter wheel, focusing lenses, and a CMOS camera. The objective lenses of the excitation and detection arms are mutually orthogonal to the sample position. A waveform generator synchronizes the galvanometer mirrors and the electrotunable lens, ensuring that every point on the oblique light sheet is well focused onto the CMOS detector. The waveform generator also triggers the camera to synchronize its line-by-line image capture (progressive mode) with the laser’s position. To generate 3D tomograms (i.e., 2D oblique image stacks; see Supplementary Figure 5 for a comparison of 3D images obtained by the flow zoometer and a confocal fluorescence microscope), the entire capillary is scanned along the flow axis by a stepper motor stage. This scanning occurs once for each channel of fluorescence, at a rate of 1 larva per 17 sec per color. The 4-color image number of voxels and data size are 6 Gigavoxels and 12 GB per animal, respectively. The lateral resolution of the flow zoometer is approximately 1 μm, constrained by the microscope magnification, while the axial resolution is typically 1-10 μm, dictated by the scan speed and width of the light sheet (see Supplementary Figure 6 for the evaluation of the lateral and axial resolutions). The robotic arm, fluidic delivery system, and LSM are coordinated via custom software. The flow zoometer can flow and image more than 1,000 larvae within 24 hr (see Supplementary Figure 2b for a time-series representation of 3D tomogram acquisitions).

The fifth component is an AI-based image processor which reconstructs 3D images and applies segmentation and classification algorithms to assess the biological condition of each larva. Initially, the acquired raw 3D images undergo an affine transformation to adjust for the oblique angle of the imaging plane, followed by downsampling. Segmentation algorithms are used to pinpoint regions of interest in the reconstructed 3D images (see Supplementary Figure 7 and Supplementary Methods 1 and 2 for details of the segmentation methods). In the final step, a Random Forest classifier is employed to categorize the internal state of each individual larva. A strength of the flow zoometer is its modularity; the components can be used independently or in concert as needed.

### Basic performance of flow zoometry

To show the basic performance of the flow zoometer, we performed flow zoometry on populations of larvae comprising several different genotypes. These included DAPI-stained^34^ (imaged in the blue channel) larvae expressing green fluorescent protein (GFP; green channel) in fat body cells, membrane-targeted tandem Tomato fluorescent protein (mtdTomato; orange channel) in wing discs, and red fluorescent protein (RFP; red channel) in the Bolwig organ, nervous system, hindgut, and anal plate (*UAS-mtdTomato,10xQUAS-6xGFP/nub-GAL4;r4-QF2^3xP3-RFP^/+* with DAPI staining; see Supplementary Note 1). Next, we imaged a strain expressing GFP (green channel) in the peripheral nervous system and ventral nerve cord (*ppk-GAL4,UAS-mCD8::GFP*)^35^. The two remaining strains were one expressing Tomato fluorescent protein (orange channel) in the central nervous system (*UAS-mCD8::Tomato;drl-GAL4*) and one expressing monomeric RFP (mRFP; red channel) in wing discs, haltere discs, salivary glands, and proventriculus (*nub-GAL4,UAS-myr::mRFP*).

Figure 2a through Figure 2d present representative orthographic projections of 3D tomograms obtained from the four genotypes, as detailed in Supplementary Video 3 which shows the tomograms in three dimensions and Supplementary Figure 8 which shows white-on-black greyscale projections. In the case of the 4-color tomograms (Figure 2a), each channel revealed fluorescence from a different exogenous protein type. Combining genetic engineering with multi-color 3D tomography, the flow zoometer highlights multiple tissues of interest against the background of other tissues. In the 1-channel tomograms of the *pickpocket(ppk)-GAL4* strain (*ppk-GAL4,UAS-mCD8:GFP*) (Figure 2b), individual GFP-expressing peripheral neurons stood out against the autofluorescence background. In the 1-channel tomograms of the *derailed(drl)-GAL4* strain (*UAS-mCD8::Tomato; drl-GAL4*) (Figure 2c), we identified the central nervous system including the brain lobes, ventral nerve cord, and segmentally organized nerves, with all components well contrasted against the autofluorescence background. In the 1-channel tomograms from the *nubbin(nub)-GAL4* flies (*nub-GAL4,UAS-myr::mRFP*) (Figure 2d), we were able to distinguish the central areas of both wing discs, the haltere discs, salivary glands, proventriculus, ventral nerve cord, and individual midgut epithelial cells. The images in the figures underscore the broad applicability of the flow zoometer; it can effectively image fluorescence from a variety of exogenous fluorescent proteins with high contrast against an autofluorescence background, across a range of colors and tissues, down to single-cell resolution (see individual denticles shown in the lower-right inset of Figure 1 and individual neurons, midgut epithelial cells, and segmentally organized nerves shown in Figure 2 and Supplementary Figure 8).

**Figure 2.**
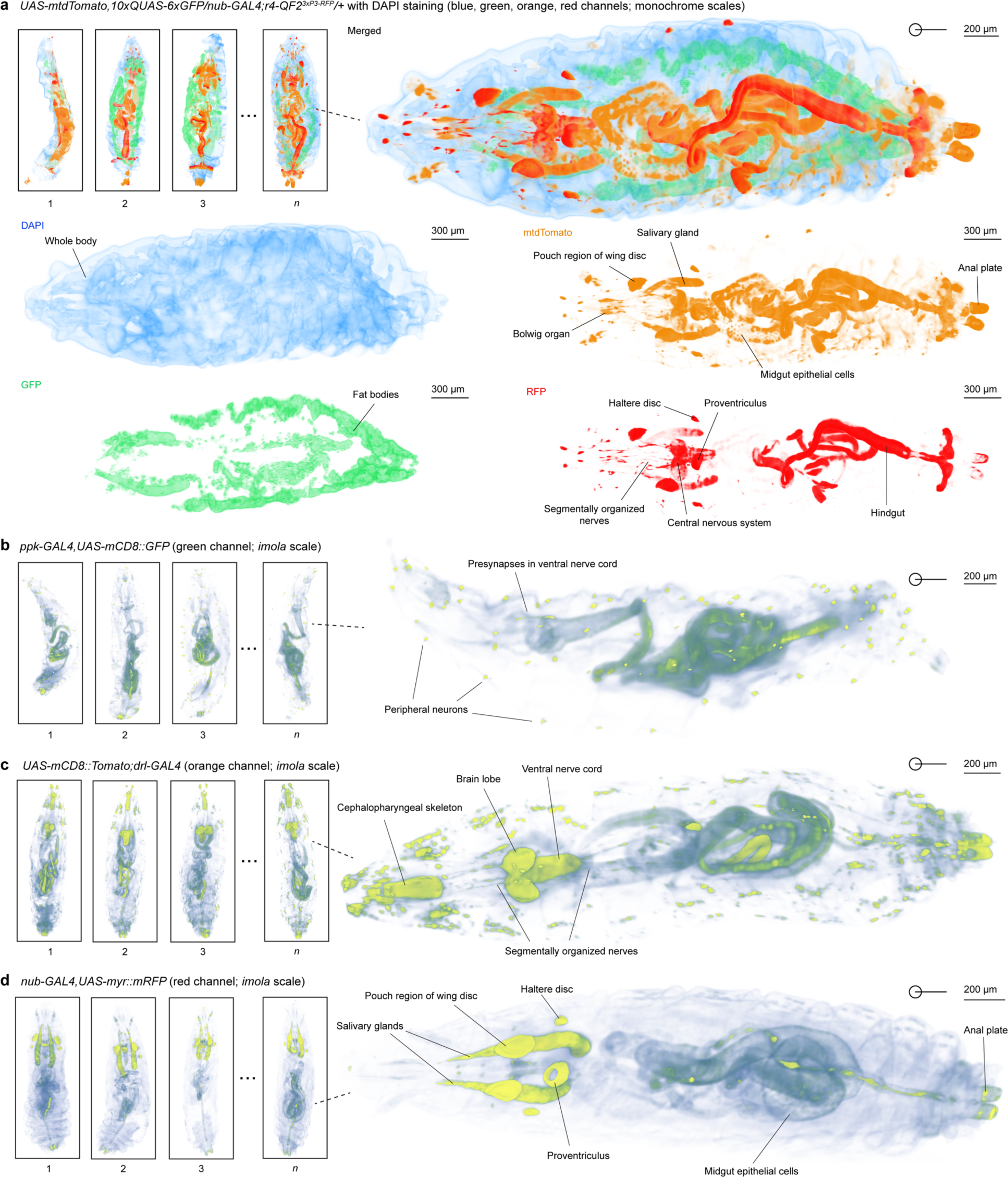
3D tomograms of various types of *Drosophila* acquired by flow zoometry. **a,** 4-channel 3D tomogram (blue, green, orange, and red channels) of 4-color larvae (*n* = 11). In the blue channel, DAPI staining enabled the visualization of the whole body. In the green channel, GFP expression revealed fat bodies throughout the larva. In the orange channel, mtdTomato was expressed in various tissues including the Bolwig organ, pouch region of wing discs, salivary glands, midgut epithelial cells, and anal plate. In the red channel, RFP expression highlighted multiple tissues including segmentally organized nerves, the central nervous system, haltere discs, and proventriculus. **b,** 1-channel tomograms (green channel) of *ppk-GAL4* flies (*n* = 37) expressing GFP in peripheral neurons. **c,** 1-channel tomograms (orange channel) of *drl-GAL4* flies (*n* = 50) expressing Tomato in the cephalopharyngeal skeleton, the central nervous system including the brain and ventral nerve cord, and the segmentally organized nerves. **d,** 1-channel tomograms (red channel) *of nub-GAL4* flies (*n* = 123) expressing mRFP in the haltere discs, pouch regions of the wing discs, salivary glands, proventriculus, midgut epithelial cells, and anal plate.

Big tomographic data of *nub-GAL4* larvae (*n* = 422; see Supplementary Figure 9 for a library of 3D tomograms obtained in a representative experiment) obtained across several more measurements revealed morphological feature distributions within a population with high statistical confidence. A 3D U-Net segmentation algorithm^36^ was employed to identify and quantify the morphology of target tissues: the central regions of wing discs, the haltere discs, salivary glands, and proventriculus (Figure 3a). Depicted in Figures 3b through Figure 3g are the results of the flow zoometer’s multidimensional statistical analysis. Shown in Figure 3b, Figure 3c, and Figure 3d are histograms of the volumes of the central regions of the left and right wing discs, the volumes of the left and right haltere discs, and the volumes of the left and right salivary glands, respectively. Our calculations identified no statistically significant left-right differences in the volumes of the salivary glands, wing discs, or haltere discs. Shown in Figure 3e and Figure 3f are 2D scatter plots of total wing disc volume and body length, and total haltere disc volume and total wing disc volume, respectively. Figure 3g shows a t-SNE plot of the inertia tensors calculated from the wing discs, haltere discs, salivary glands, and proventriculus. The inertia tensors of each class form separate clusters, which may indicate that each class has intrinsic differences in shape, size, directionality, or density distribution. These differences in the characteristics of the tissues are captured by the segmentation algorithm, representing a critical step for further data interpretation and method application.

**Figure 3.**
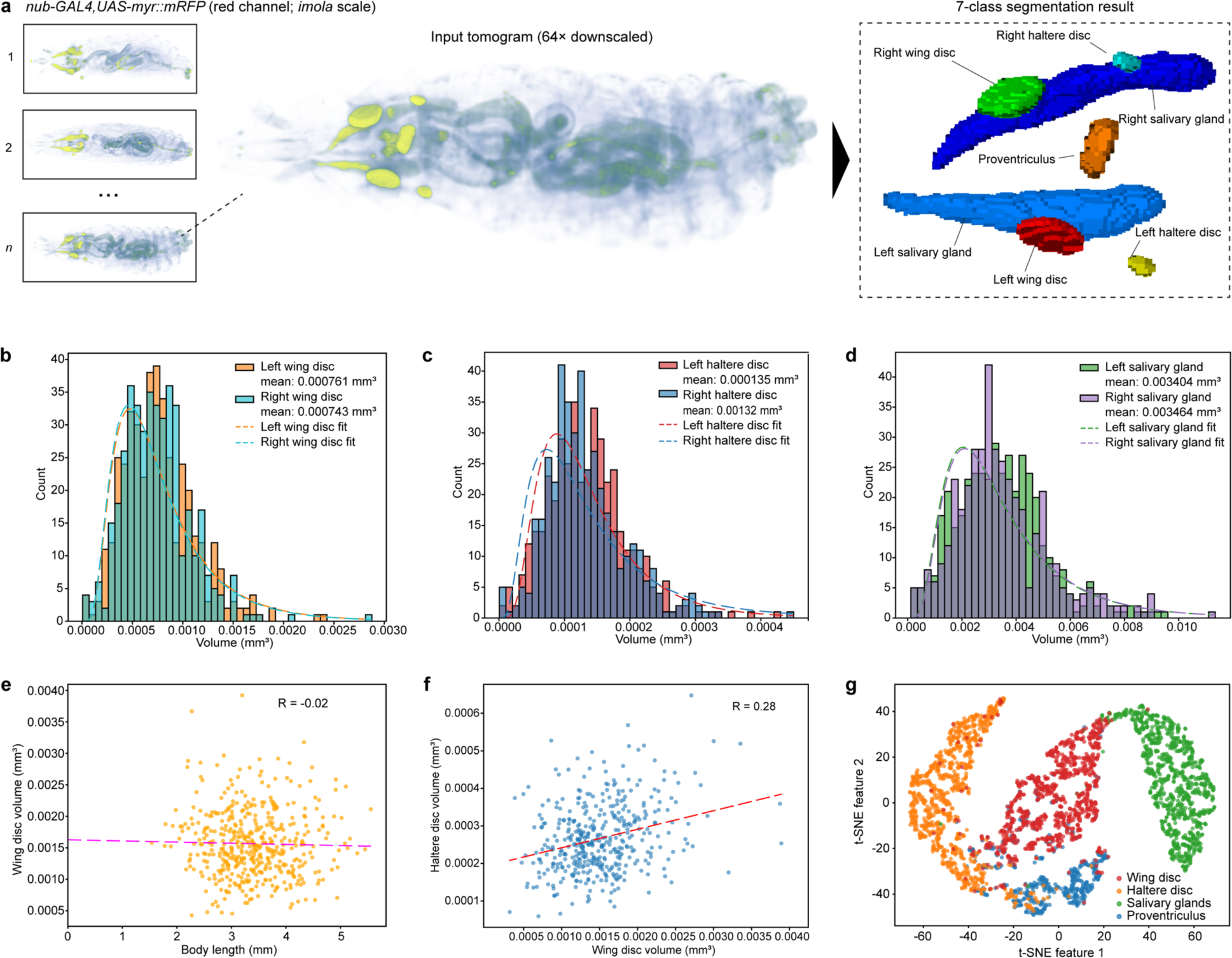
Data-driven analysis of *Drosophila*. **a,** Principles of our data-driven analysis of *Drosophila* including the segmentation of 7 tissue classes from the tomogram of a whole *Drosophila* larva. In the obtained whole-body tomogram (left), multiple fluorescence expression patterns, including those originating from fluorescent proteins and autofluorescence, were detected. By applying the developed 3D U-Net segmentation algorithm, each of 7 tissue classes was isolated from the whole-body image and used for downstream statistical analysis. **b,** Histograms of central region volumes for left (orange trace) and right (blue trace) wing discs. Standard deviations (SDs) of left and right wing disc volumes are 3.55 × 10^-3^ mm^3^ and 3.29 × 10^-3^ mm^3^, respectively. **c**, Histograms of the volumes of the left (red trace) and right (blue trace) haltere discs. SDs of left and right haltere disc volumes are 6.13 × 10^-6^ mm^3^ and 6.05 × 10^-6^ mm^3^, respectively. **d,** Histograms of the volumes of the left (green trace) and right (purple trace) salivary glands. SDs of left and right salivary gland volumes are 0.00163 mm^3^ and 0.00174 mm^3^, respectively. **e,** 2D scatter plot of total wing disc volume and body length. **f,** 2D scatter plot of total haltere disc volume and body length. **g,** t-SNE plot of the inertia tensors calculated from the 7 segmented tissues. Left and right classes were combined into a single class for the wing discs, haltere discs, and salivary glands. All plots were calculated from the same big tomographic data of *nub-GAL4* flies (*n* = 422). The fits in **b-d** are log-normal, while those in **e** and **f** are linear.

### Application to large-scale whole-animal toxicological screening

To highlight the utility of our flow zoometer in toxicology and food science, we performed a quantitative analysis of phenotypic changes in *Drosophila* larvae exposed to environmental microplastics (MPs), the effect of which is not well understood during organism development. Although whole-animal toxicology assays typically require skilled manual handling and analysis of costly vertebrates^37^, flow zoometry of *Drosophila* provides a labor-free, affordable option. As illustrated in Figure 4a, we first mixed various MPs into food upon which adult *Drosophila* laid their eggs. The resulting larvae ate the MP-infused food during their development and were subsequently analyzed by flow zoometry. Larval body length, body volume, and wing disc volume were segmented using the 3D U-Net algorithm and quantified. We tested two concentrations (0.25 µg/g food and 2.5 µg/g food, or 0.26 µg/g food and 2.6 µg/g food) each of 500 nm and 4 µm polystyrene (PS), 15 µm poly(methyl methacrylate) (PMMA), and 6 µm polytetrafluoroethylene (PTFE) particles. Control samples of *Drosophila* not exposed to MPs were prepared and analyzed in tandem for each plastic type. In total, we tissue-cleared and analyzed 1,352 larvae in 110 hr (Supplementary Table 1).

**Figure 4.**
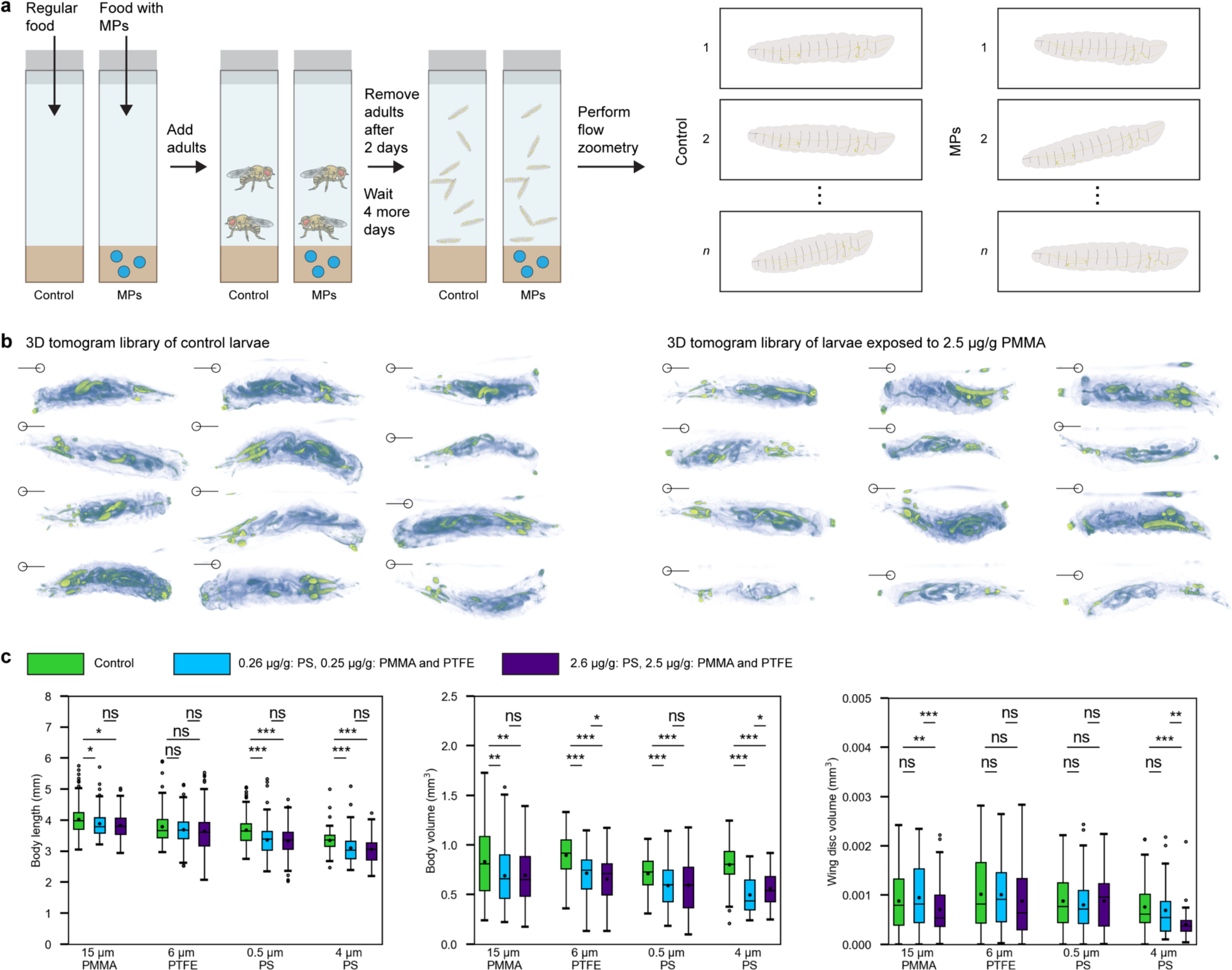
Application to large-scale whole-animal toxicological screening. **a,** Schematic of the experiment. Regular food (control) and food with various types and concentrations of microplastics (MPs) were prepared. *Drosophila* adults were added to the food vials and laid eggs. The resulting larvae consumed the regular or MP-infused food, grew to the L3 stage, and were collected and analyzed by flow zoometry. **b,** 3D tomogram library of control larvae (left) and larvae exposed to 15 µm PMMA MPs (2.5 µg/g food). **c,** Box plots of *Drosophila* body length (*n_1_* = 1,337), body volume (*n_2_* = 1,274), and wing disc volume (*n_3_* = 1,520) calculated by our 3D U-Net segmentation algorithm for the four types of MPs. For each MP type, we tested two loading amounts and a control sample. Plotted (*n_1_* = number of body lengths; *n_2_* = number of body volumes; *n_3_* = number of wing discs) are: 15 µm PMMA control (*n_1_* = 149; *n_2_* = 141; *n_3_* = 190), 0.25 µg/g (*n_1_* = 129; *n_2_* = 114; *n_3_* = 162), and 2.5 µg/g (*n_1_* = 119; *n_2_* = 109; *n_3_* = 156); 6 µm PTFE control (*n_1_* = 108; *n_2_* = 105; *n_3_* = 102), 0.25 µg/g (*n_1_* = 106; *n_2_* = 104; *n_3_* = 125), and 2.5 µg/g (*n_1_* = 120; *n_2_* = 114; *n_3_* = 116); 0.5 µm PS control (*n_1_* = 110; *n_2_* = 110; *n_3_* = 156), 0.26 µg/g (*n_1_* = 112; *n_2_* = 109; *n_3_* = 148), and 2.6 µg/g (*n_1_* = 121; *n_2_* = 117; *n_3_* = 139); and 4 µm PS control (*n_1_* = 73; *n_2_* = 71; *n_3_* = 114), 0.26 µg/g (*n_1_* = 107; *n_2_* = 102; *n_3_* = 46), and 2.6 µg/g (*n_1_* = 83; *n_2_* = 80; *n_3_* = 66). Results of unpaired *t* test (two-sided) are shown between plot pairs. A *p* value of over 0.05 is designated as “ns”; *p* < 0.05 is “*”; *p* < 0.01 is “**”; and *p* < 0.001 is “***”. Box limits: interquartile range (IQR); whiskers: 1.5 × IQR beyond box limit; points outside whiskers: outliers; line in box: median; dot in box: mean.

Shown in Figure 4b are representative 3D tomograms of control larvae and larvae exposed to 2.5 µg/g PMMA. Persisting morphological differences between the two cohorts are not evident against the background of individual-level variation. Shown in Figure 4c are box plots of the body length, body volume, and wing disc volume for each MP type and its control sample (see Supplementary Table 2 for detailed results). As shown in Figure 4c, the calculated mean body lengths of 3 of 4 samples exposed to MPs were significantly lower than those of their corresponding control samples (see Supplementary Figure 10 for Monte Carlo simulations of sample size on statistical significance in this experiment). This result is in accordance with that of a previously reported small-scale study reporting *Drosophila* larval body length change due to exposure to PS particles in food^38^. Likewise, the calculated mean body volumes of all samples exposed to MPs were lower than those of their corresponding control samples. These results indicate that food-based MP exposure during the early stages of the *Drosophila* life cycle may interfere with normal body development. We also quantified total wing disc volume in each cohort. While we found no statistically significant differences between the control and lower-MP-exposure cohorts, we found statistically significant reductions in wing disc volume between the control and higher-MP-exposure 15 µm PMMA and 4 µm PS samples, indicating that the development of wing discs may be less affected by food-based MP exposure than the body volume and length, and that wing disc development may be affected by exposure to certain types of MPs but not others. Hence, we demonstrate the comprehensive quantification of body volume and internal tissue volume changes in *Drosophila* as a result of their exposure to MPs, all within only 5 days by flow zoometry.

### Application to large-scale whole-animal genotypic screening

To demonstrate the effectiveness of our flow zoometer in genetic research, we automated the widely-used genotypic screening process for individual *Drosophila*. This allowed us to perform high-resolution 3D tomographic imaging only of the targeted individuals, resulting in considerable time and effort savings compared to traditional manual techniques used in *Drosophila*-based genetic research, where target individuals must be isolated one by one by skilled hand and identified by phenotypic signs^17^. Figure 5a demonstrates how the flow zoometer facilitated this process on-line by conducting rapid low-resolution scans automatically on each individual to determine the presence of (i) a Tubby phenotype derived from a balancer chromosome (*CyO-Tb-RFP*) which enables morphology-based genotyping or (ii) mCherry or EGFP fluorescence which enables fluorescence-based genotyping. We then obtained high-resolution images of only the target flies, while the non-target larvae (those carrying the balancer chromosome that are not required for subsequent analysis) were released without further measurement.

**Figure 5.**
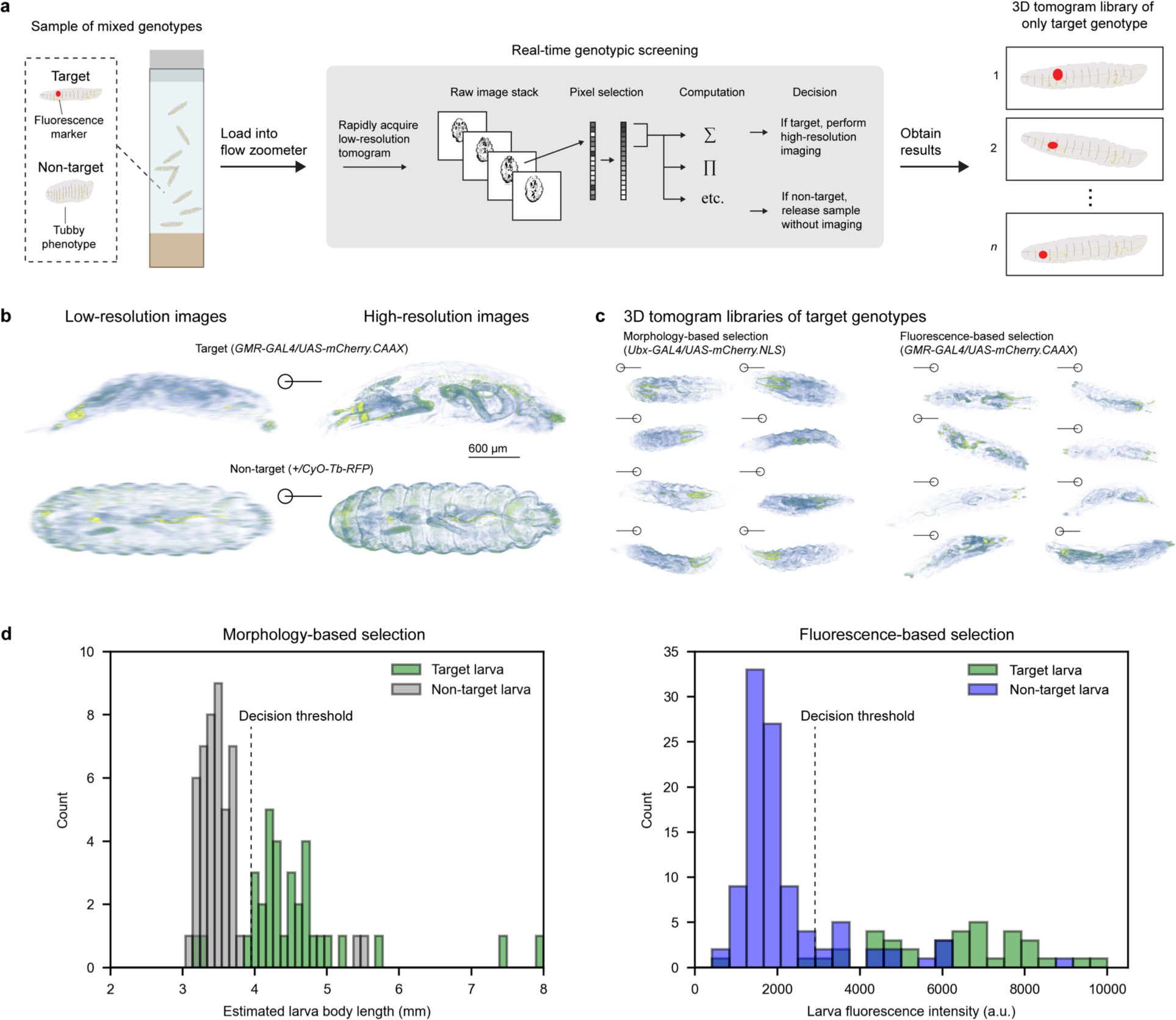
Application to large-scale whole-animal genotypic screening. **a,** Schematic of the experiment. A sample of mixed genotypes containing target and non-target samples was loaded into the flow zoometer. Each larva was measured rapidly. The flow zoometer performed automated quantitative analysis on the raw image data. Pixel subsets (e.g., highest-intensity *n* pixels from each of *m* frames) were identified from the image stack, and relevant computations (e.g., arithmetic mean) were performed on the pixel subsets. The results of the computations were screened against threshold criteria to identify the sample genotype via its body phenotype or tissue fluorescence. Target samples were subsequently imaged at high resolution, while non-target samples were released. **b,** Representative 1-channel 3D tomograms of target (upper) and non-target (lower) larvae taken by flow zoometry in the rapid imaging (left) and high-resolution (right) imaging modes. The high scan speed resulted in low axial resolution in the reconstructed 3D tomograms, while the high-resolution tomograms revealed far finer phenotypic details. 3D tomograms are shown in scientific *imola* color scale. **c,** 3D tomogram libraries of high-resolution images of target larvae from the morphology-based and fluorescence-based genotypic screening. **d,** Histograms of target (*n* = 34) and non-target (*n* = 46) samples measured in body length-based selection (left plot; bin width 0.1 mm) and target (*n* = 37) and non-target (*n* = 100) samples measured in fluorescence-based selection (right plot; bin width 400 a.u.). Also plotted are the decision thresholds used for each genotypic screening experiment.

Figure 5b presents representative rapid-scan tomograms of target and non-target flies. Despite the lower resolution, the flow zoometer was able to differentiate between target and non-target flies via two methods for automated quantitative analysis. Each low-resolution tomogram was acquired in 7 sec and analyzed in less than 4 sec. We performed morphology-based real-time genotypic screening (see Methods for more details) during flow zoometry of a sample of 100 larvae (50 target, 50 non-target). The flow zoometer investigated the genotypes of 80 larvae (20 lost samples) and obtained 33 high-resolution tomograms (representative 3D tomograms shown in Figure 5c, left). Within the 33 high-resolution tomograms, we manually identified 31 targets, corresponding to a genotyping purity and yield of 94% and 62%, respectively. From the loading of the larvae to the acquisition of the final tomogram, the entire experiment took only 112 min. To demonstrate fluorescence-based genotypic screening, we loaded 100 larvae (50 target, 50 non-target) and screened 90 larvae (10 lost samples). Within the 30 obtained high-resolution tomograms (Figure 5c, right), we identified 30 targets, corresponding to a genotyping purity and yield of 100% and 60%, respectively. Histograms plotting the distributions of single-genotype samples (manually prepared) screened by both methods are shown in Figure 5d. By automating genotypic screening during flow zoometry, we managed to identify and image specific individuals with target genes with high accuracy, thereby circumventing the need for labor-intensive manual genotyping and lengthy data acquisition.

### Application to large-scale whole-animal tumor screening

To showcase the capabilities of our flow zoometer in tumor screening, we evaluated its utility in identifying the cellular transformations caused by genetic mutations in a *Drosophila* model of pancreatic ductal adenocarcinoma (PDAC). Given its 5-year relative survival rate of only 13%^39^, PDAC is one of the most lethal cancer types, highlighting the needs for both mechanistic and therapeutic studies. Establishing an automated method to quantify the generation and progression of transformed cells in a PDAC model at the individual organ level could significantly advance our understanding of carcinogenesis, including the pathophysiological interactions between transformed cells and the host. It could also accelerate the analysis of individual differences and the identification of potential therapeutic targets effectively.

Our previous studies using a ‘*4-hit*’ *Drosophila* model mimicking the PDAC genotype (alterations of the oncogene *KRAS* and tumor suppressor genes *TP53*, *CDKN2A*, and *SMAD4*)^40,41^ required labor-intensive, visually-based viability assays to monitor disease progression, a method that only offers a binary assessment of disease presence and may overlook finer details of transformation in each animal. To address these limitations, we refined the model to *ptc-GAL4,UAS-mCherry,UAS-4-hit*, in which transformed cells are labelled with *patched*(*ptc*)*-GAL4*-driven mCherry which offers a lower autofluorescence background and deeper tissue penetration, allowing for more efficient 3D tomography than GFP^40–42^. We performed large-scale flow zoometry on both control without the transformation-causing transgenes [*ptc-GAL4, UAS-mCherry* (abbreviated as *ptc>mCherry*); *n* = 1,309] and tumorigenic [*ptc-GAL4, UAS-mCherry, UAS-4-hit* (abbreviated as *ptc>mCherry 4-hit*); *n* = 1,450] genotypes and employed a Random Forest classifier to distinguish tumorigenic from non-tumorigenic larvae on a continuous probability scale.

Figure 6a depicts a schematic of our tumor screening. Figure 6b displays confocal fluorescence microscopy images of dissected wing discs (shown in white outline) from both control (*ptc>mCherry*; left) and tumorigenic (*ptc>mCherry 4-hit*; right) larvae. Owing to the proliferation and migration of transformed cells, the morphology of the tumorigenic *ptc* regions were significantly different than those of their non-tumorigenic counterparts, similar to our previous observations^40,41^, validating the developed PDAC model. Figure 6c shows 3D tomograms of the *ptc* regions of the wing discs of the control (*ptc>mCherry*; left) and tumorigenic (*ptc>mCherry 4-hit*; right) larvae obtained in whole animals via flow zoometry separately from the large-scale screening experiment. The phenotypic differences between the control and tumorigenic *ptc* regions observed *in toto* via flow zoometry are consistent with those in the dissected wing discs. In Figure 6d are representative 3D tomograms and resulting segmentation masks from the large-scale screening of control (upper; *n* = 937) and tumorigenic (lower; *n* = 850) larvae used for the automated tumor screening experiment. Prediction scores of the ground truth are shown to the right of the masks. The top four masks of each type represent the best predictions, while the bottom two represent the worst predictions. In general, the *ptc* regions in successfully predicted control samples follow the well-defined shape shown in Figures 6b and 6c, while those in successfully predicted tumorigenic samples are more diverse in their morphological characteristics, matching what is seen in Figures 6b and 6c. Figure 6e compares the screening performance across multiple experiments (k-replicate nested cross validation as explained in Methods) through receiver operating characteristic (ROC) curves, showcasing an average accuracy 67.87% with a standard deviation of 5.42% (see Supplementary Method 2 for details). These results underscore the potential of flow zoometry for precise, automated quantification of tumor progression, offering promising avenues for future applications in cancer biology and pharmacology.

**Figure 6.**
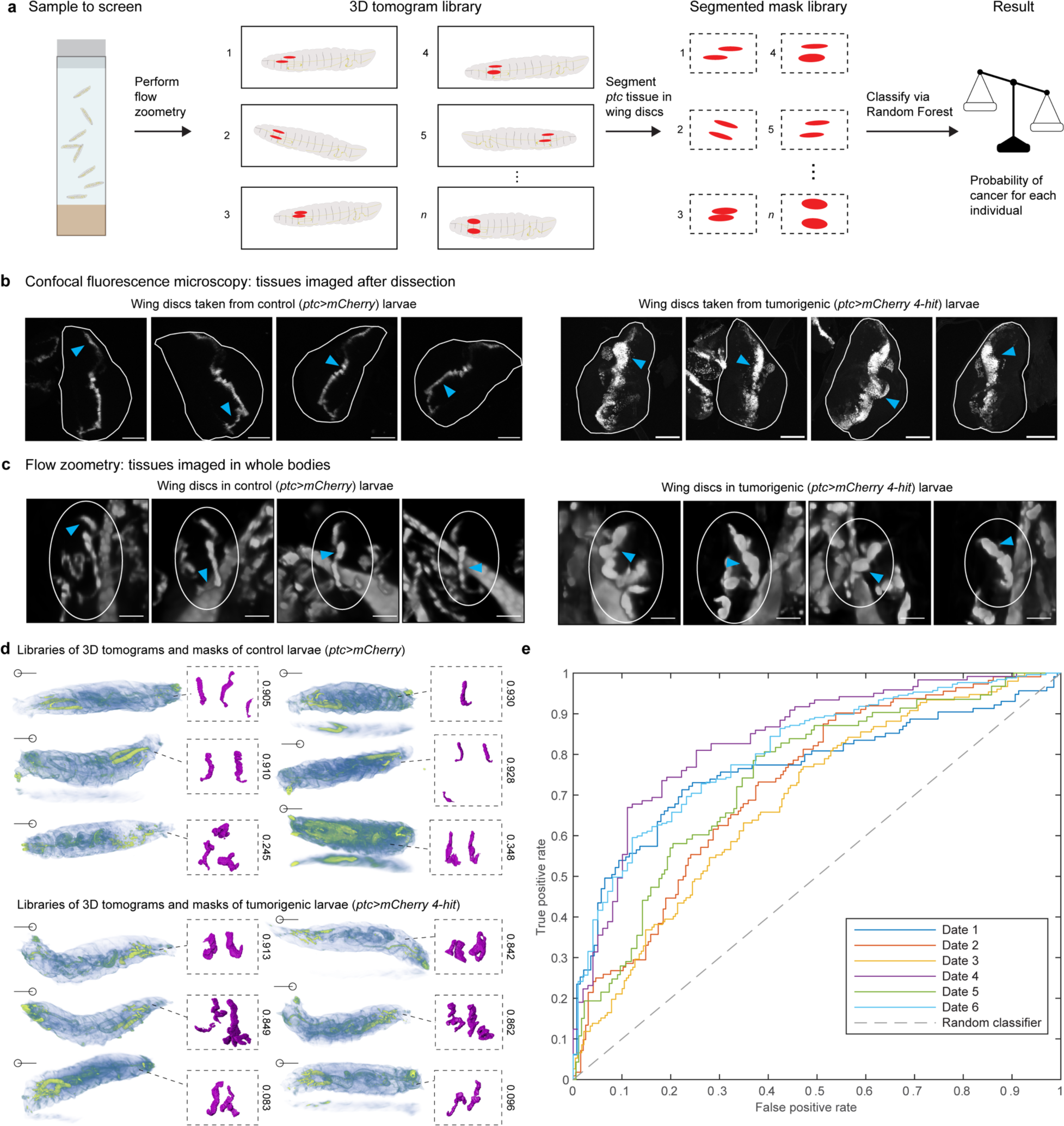
Application to large-scale whole-animal tumor screening. **a,** Schematic of automated tumor screening. A 3D tomogram library is obtained, and the *ptc* tissue from wing discs is segmented. The segmented masks are classified by the Random Forest classifier. **b,** Confocal microscopy images of dissected wing discs (shown in white outline) taken from control (*ptc>mCherry*; left) and tumorigenic (*ptc>mCherry 4-hit*; right) larvae. Scale bars: 100 µm. **c,** Representative 3D tomograms (red channel) of control *ptc>mCherry* (left) and tumorigenic *ptc>mCherry 4-hit* (right) larvae obtained by flow zoometry. Cellular transformation in the *ptc>mCherry 4-hit* larvae causes noticeable change in the morphology of the *ptc* region of wing discs in both confocal fluorescence microscopy and flow zoometry images (arrows). Tomograms are rotated in 3D space such that the orientations of the *ptc* regions approximately match those in **b.** Scale bars: 100 µm. **d,** Representative 3D tomograms and their resulting segmentation masks from the big tomographic data of control *ptc>mCherry* (upper; *n* = 937) and tumorigenic *ptc>mCherry 4-hit* (lower; *n* = 850) larvae used for automated tumor screening. Tomograms are plotted with the same intensity and size scales and orientation to the projection plane. Masks are rotated in 3D space to approximately match the orientation of the tomograms in **b** and **c**. **e,** Random Forest classification results shown as calculated receiver operating characteristic (ROC) curves obtained across multiple experiments.

## DISCUSSION

Flow zoometry is a transformative technology, reshaping the scope of biological research in *Drosophila* by facilitating a fully automated, whole-body analysis at a scale and precision previously unachieved. This advancement propels the utility of *Drosophila* as a model organism, providing an exhaustive level of analysis that surpasses what was possible with research confined to embryonic stages. By automating the process, flow zoometry not only alleviates the burden of manual labor but also uncovers organism-level structural changes that are generally overlooked in conventional morphological analysis of *Drosophila* that relies on dissecting, separating, and imaging individual larval structures. Automated assay systems have demonstrated their power in animal behavioral research to obtain such unexpected findings from large sample sizes (e.g., a rare sleepless population in adult *Drosophila*^43^). Analogously for morphological studies, flow zoometry’s ability to detect subtle effects or rare phenomena, otherwise obscured by variations at the individual level, is a testament to its potential. Our extensive, whole-animal, high-content screening experiments have demonstrated the aforementioned advantages of flow zoometry. Analysis from the experiments revealed considerable variability in the size of bodies and organs across stage-matched L3 larvae. This heterogeneity underscores the importance of analyzing a broad array of animals per genotype and assessing multiple parameters per individual. We found a correlation between the volumes of wing discs and haltere discs, indicating their concurrent development during the L3 stage. However, our studies found minimal correlation between the volume of imaginal discs and the overall body volume, suggesting a disconnect in their developmental timelines through the L3 stage. Notably, our investigations did not identify any significant left-right asymmetries in the volume of wing discs, haltere discs, or salivary glands, while a prevailing left-favored asymmetry in adult fly wings has been previously reported^44^. Our flow zoometry data suggests that the documented asymmetries in adult *Drosophila*, such as the larger-left-wing phenomenon, may manifest later than at the L3 stage, potentially during pupation.

This technology notably enriches the morphological data we can gather per animal, enhancing our ability to spot subtle deviations from the norm associated with experimental manipulations (Supplementary Figure 10). For instance, our toxicology studies using flow zoometry to examine the impact of MP exposure on larvae revealed statistically significant effects on their body length, body volume, and wing disc volume. We also identified differences between tissue sensitivity to MP exposure judged by the effects of MP exposure on tissue size. Our flow zoometer also performed automated segmentation and classification of transforming internal tissues in our model for PDAC, facilitating the swift screening of tumorigenicity. These diverse and promising results underscore both the breadth and power of flow zoometry of *Drosophila* in biological research applications, of which we believe our demonstrations are not exhaustive.

Looking forward, flow zoometry stands out as a promising tool for investigating morphological phenomena or physical responses across multiple organs while their orientation and connectivity in each animal remain unperturbed. This aligns well with current trends in biological research, such as the intricate connections between brain neurons and digestive organs^45^ or the systemic actions of hormones derived from neurosecretory cells orchestrating metabolism and behavior^46^. By preserving the intact structure of organisms, flow zoometry facilitates comprehensive studies of the spatial relationships of cells in tissues and their interrelationships, emerging as a new “spatial biology” field. Furthermore, flow zoometry is suited for studying rare events which require large sample sizes to ensure statistical significance. For example, it could advance our understanding of “loss of heterozygosity” caused by mitotic crossover events in somatic tissue, which is known to occur at very low frequencies of ∼0.01% in some tissues^47^. Another potentially powerful application of flow zoometry is in mapping whole-animal distributions in a manner analogous to cell-type mapping in a given cell population via flow cytometry. By employing multiple markers to identify various disease-associated phenotypes, organ functions, and other abnormalities in *Drosophila*, flow zoometry could be used to map whole-animal phenotypic or symptomatic variations. This, in turn, would enable the cataloging of physiological and disease-related organ-organ interactomes, offering new insights into the dynamics of health and disease at the organism level.

The flow zoometry system we have developed has a number of promising areas for enhancement and extension (see also Supplementary Discussion 1). First, although we currently rely on the fixation of larvae, flow zoometry of live *Drosophila* larvae is possible in principle. By leveraging emerging advances in fluorescence microscopy in the infrared region^48^, such as multiphoton imaging^49^ or specialized near-infrared probes^50^, whole-body 3D imaging throughout un-cleared living larvae may be possible. To avoid motion blur in obtained tomograms, non-lethal anesthesia offers a solution: reported anesthesias^51^ are effective on time scales of hours and could be administered using our automated tissue-clearing device. Flow zoometry of live *Drosophila* larvae could be combined with individual-level sorting to enable valuable downstream assays including directed evolution or time-based observation spanning multiple life cycle stages. Second, while the current system was designed for and is limited to *Drosophila* at life cycle stages preceding the pupa stage, it should be possible in principle to modify the system to accommodate *Drosophila* at the adult stage. Tissue clearing and 3D imaging of adult *Drosophila*^32^ are likely possible in our current system, while the handling of adults with fragile appendages could be achieved by redesigning the fluidic delivery system. Large-scale flow zoometry of adult *Drosophila* would enable statistically significant analysis of the phenotypes of anatomical systems including fully developed wings and eyes, both model tissues routinely scrutinized to genetically elucidate interactions between genes as well as signaling pathways^1,10,11,13,22^. Third, the incorporation of super-resolution fluorescence imaging techniques to enhance the lateral and axial imaging resolutions and achieve sub-cellular volumetric imaging^52^ in flow zoometry would enable large-scale statistical analysis of small targets such as cell mitochondria or nuclei. Considering the resulting large data size, it may be necessary to also implement a real-time AI system which takes super-resolution tomograms of only target regions. Such an algorithm may resemble our genetic screening algorithm which makes decisions based on low-resolution tomograms. Looking toward the future, by implementing robotic husbandry^53^ and reinforcement learning algorithms which can direct experiments^54^, flow zoometry might be expanded to realize a completely self-contained and autonomous system for *Drosophila* genetic design, breeding, analysis, and experimental exploration.

In summary, flow zoometry heralds a major leap in high-throughput screening technology, extending its capabilities to encompass comprehensive studies of entire *Drosophila*. It not only elevates the precision of biological research but also opens the door to serendipitous discoveries across various areas of biology. Flow zoometry is a pivotal “AI for science” platform,^55^ a new standard which enables detailed examination of whole animals and promises to accelerate advancements in genetics, developmental biology, neuroscience, and more.

## METHODS

### Tissue-clearing device

The core compartment of the tissue-clearing device, as depicted in Supplementary Figure 1a, was fabricated from polypropylene (PP) using 3D printers (Prusa i3 MK3S+, Qidi X-Plus II). The PP material was chosen due to its resilience to common tissue-clearing reagents, including those employed in our protocol. The chamber design incorporated 6 silicone inlet tubes for reagent dispensation, 6 sample vials for larval containment, and an outlet tube. The inlet tubes were linked to reagent reservoirs, and the required reagents were drawn into the main chamber using peristaltic pumps at a rate of 25 mL/min. The sample vials consisted of 15-mL PP centrifuge tubes, modified with polyethylene mesh screens (∼400 µm holes) replacing their conical bottoms. The outlet tube was designed to split into multiple channels, selected by solenoid pinch valve and peristaltic pump, to ensure appropriate separation of waste types. Control over the peristaltic pumps and pinch valves was achieved using a custom power supply switcher and controller (Arduino Mega 2560), allowing precise management of administered reagent volumes and exposure times.

### Tissue-clearing protocols

Larvae in food tubes were washed out with distilled water, and the larvae were strained out. The larvae were then washed with a 4% bleach solution (Wako 197-02206) for 10 min for optional dechorionation. For automated fixation, live larvae were loaded into the tissue-clearing device and initially rinsed 3 times with distilled water for 5 min each time. Following this, they were fixed with a 4% paraformaldehyde in a 0.1 M phosphate buffer solution (4% PFA in PBS; Wako 163-20145) for a duration of 2 hr at room temperature. For delipidation, larvae were exposed to 0.2% Triton X-100 (Wako 169-21105) in 0.1 M PBS (PBT) 3 times for 90 min each. Afterward, the larvae were exposed to escalating concentrations of 1-propanol (Wako 162-13461) in distilled water for dehydration, which involved 120-min exposures to each of the following concentrations: 10%, 25%, 50%, 60%, 80%, and 99.7% (separated into 2 × 60-min exposures) 1-propanol. All dehydration steps, with the exception of the final 99.7% 1-propanol phase, were conducted at pH 9, which was adjusted using triethylamine. Post-dehydration, the larvae were exposed to ECi for more than 12 hr prior to imaging. For the genotypic screening, tumor screening, and 4-color and *drl-GAL4* larvae in the proof-of-concept experiments, the stages of dechorionation, fixation, and delipidation were manually conducted at Hokkaido University and Tohoku University using the same reagents as those used at the University of Tokyo. Instead of the 2-hr exposure to 4% (9% for the *ppk-GAL4* larva) PFA in PBS at room temperature, the step was conducted overnight at 4 °C. After the final delipidation step with PBT, larvae were placed into PBS and transported to the University of Tokyo, arriving within 2-3 days. Upon arrival, the larvae were inserted into the tissue clearing device, and the clearing protocol continued in a fully automated fashion starting from the dehydration stage. The division of the tissue-clearing process across two labs was a necessary step to adhere to Japanese regulations concerning the handling of genetically modified organisms. We found no differences between wild-type or *nub-GAL4* larvae that underwent the entire process within our device compared to those that experienced the divided protocol. Refer to Supplementary Table 3 and Supplementary Table 4 for details on the 2 protocols.

### Robotic sample loader

The movements of the robotic arm were programmed using Python and enacted through the arm’s proprietary software (Dobot Studio). The movement sequence was triggered by digital voltage (TTL) signals. These TTL signals originated either from the tissue-clearing device, for the movement of the first sample vial, or from the main computer that controlled both the fluidic delivery system and the LSM, for movements of subsequent sample vials.

### Fluidic delivery system

When loading larvae into the dispersing chamber of the fluidic delivery system, excess ECi was drained from the chamber via an ECi overflow relief system, which included a silicone outlet tube and a pinch valve, which was closed when larvae were flowed toward the LSM. The dispersing jet and drive flows had flow rates of 50 mL/min and 10-20 mL/min, respectively. The pickup tube position was precisely controlled by a 2-axis motorized stage. The height of the tube was gradually lowered throughout experiments such that the rate of picking up larvae was kept relatively steady despite the reduction of their number. The fluidic chip was 3D-printed (Elegoo Saturn 3 Ultra) from resin and washed (Elegoo Mercury X). Solenoid pinch valves mounted on the imaging translation stage were placed immediately before and after the capillary, such that larvae were immobilized during imaging. The total flow rate within the capillary at the location of the microscope was approximately 70 mL/min, corresponding to a larval travel speed of about 0.029 m/s. An automated reverse-flow anti-clog system was introduced via a tubing junction following the capillary. When clogs were identified, the anti-clog system reversed the flow (200 mL/min) of the entire fluidic delivery system (ECi was drained into the ECi overflow relief system) such that blocked passages were cleared, with the clogged larvae returned to the dispersing chamber. We used a homemade motion tracker to identify (1) the presence of larvae in the dispersing chamber and (2) if larvae were exiting the dispersing chamber via the pickup tube. The motion tracker consisted of a web cam (1080P @ 30 fps) whose video data was analyzed for larval motion in the flow zoometer control software. If no motion was detected for 60 sec, the anti-clog system was engaged. If the process repeated 3 times, the next sample tube was loaded via the robotic sample loader, or the experiment was terminated. Imaged larvae were safely deposited in a mesh catcher for storage or other assays. ECi was recycled back to the ECi reservoir via peristaltic pump. Additional details of the fluidic delivery system can be found in Supplementary Figure 1b.

### Large-volume LSM

We acquired 3D tomograms with the fluorescence excitation / long-pass filter wavelengths set at 405 nm / 450 nm (bandpass filter) (blue channel), 488 nm / 558 nm (green channel), 532 nm / 600 nm (orange channel), and 594 nm / 655 nm (red channel). The *x’y’* oblique imaging plane was 2.4 mm (*x’*) × 2.7 mm (*y’*). The light-sheet size was 2 mm (*x’*) × 2.7 mm (*y’*) with a height (*y*) of 1.2 mm. In the center of the capillary, the Gaussian beam of the light sheet had a measured width (Thorlabs BP209-VIS) ∼1 µm, while the Bessel beam measured ∼2 µm, depending on alignment. Near the edge of the capillary, the Gaussian beam had a width of >5 µm, while the Bessel beam was ∼3 µm. While the Bessel beam provided better axial resolution near the edges of the capillary, we used the Gaussian beam in all experiments due to the long-term stability and beam quality reproducibility of the Gaussian beam. When imaging fluorescent beads (Polysciences, Inc. Fluoresbrite PS microspheres) to assess the LSM spatial resolution, the stage was moved at a translation speed of 0.02 mm/sec. The CMOS camera (Hamamatsu Orca Flash 4.0 V2) captured images at a rate of 20 frame/sec, corresponding to an axial resolution of 1 µm/frame. During the genotypic screening’s low-resolution scans, the stage moved at a translation speed of 5 mm/sec, which equates to 250 µm/frame. For imaging *ppk-GAL4* flies, the stage moved at a translation speed of 0.12 mm/sec or 6 µm/frame. For all other experiments, the stage moved at a translation speed of 0.2 mm/sec or 10 µm/frame, which we found was sufficient for our needs. The LSM’s magnification was set to 5.6, which yielded a lateral image resolution of 1.35 µm^2^/px. The CMOS camera captured 16-bit images measuring 2048 × 2048 pixels with an exposure time of 0.019565 msec/line. The camera was externally triggered at a rate of 20 Hz. Images were streamed over a 10 Gb/sec network and saved on an NVMe M.2 solid state hard drive in the intelligent image processor using the LabVIEW VIs provided by the camera manufacturer. Additional details of the optical setup can be found in Supplementary Figure 2.

### Intelligent image processor

Raw 3D tomographic data were read from the saved DCIMG files using the LabVIEW VIs provided by the camera manufacturer. Within LabVIEW, 2D image frames extracted from the DCIMG files were saved as HDF5 files. The raw HDF5 image stacks were affine transformed to correct for the oblique imaging plane and optionally downscaled using Python. We typically used voxel sizes of 2 µm × 2 µm × 2 µm for the reconstruction. To display each 4-color image channel, we used a linear colormap with endpoints at the figure background color (e.g., white) and a second color (e.g., blue). To display 1-channel tomograms, we used the scientific “imola” colormap^56^. The tomograms shown in Figures 2, 5c, and 6d were not downscaled. For the 3D U-Net segmentation development, quantitative analysis of *nub-GAL4* flies, *ptc* tissue segmentation and Random Forest classifier development, and tumorigenic screening experiment we performed 64× downscaling.

### 3D U-Net segmentation algorithm and nub-GAL4 larvae analysis

For analysis of *nub-GAL4* larvae, the 3D U-Net segmentation algorithm was trained using 98 1-channel (red channel) annotated tomograms of *nub-GAL4* larvae. Annotation of 7 classes of tissues (right salivary gland, left salivary gland, right wing disc, left wing disc, right haltere disc, left haltere disc, proventriculus) was performed manually in Amira 3D 2022.2 using the built-in Segmentation feature. We obtained big tomographic data of 1,380 images across 14 experiments and checked them manually, assigning quality labels (low, acceptable, and excellent quality) with a bias toward low quality. Of the total data, 9% were failed acquisitions (e.g., images of bubbles, empty images, partially-imaged larvae; *n* = 122), 24% were of low quality (e.g., larva severely out of focus, tissue was not successfully cleared, we could not clearly identify all target tissue classes; *n* = 327), 18% were of acceptable quality (we could identify the target tissues overall, but the images or certain tissues were not well focused; *n* = 254), and 49% were of excellent quality (target tissues were obviously clear and well defined against surrounding background tissues; *n* = 677). We used only the 677 excellent data for training (*n* = 98) and analysis (*n* = 579) by our 3D U-Net segmentation algorithm. For the model training, we performed preprocessing with a fixed-range Otsu threshold to remove the background, retaining only the largest connected domains and eliminating debris and reflections. The input images were first cropped to non-zero regions to focus on relevant tissue structures. Intensity normalization and resampling were then performed to normalize the input images and segment them into 80 × 112 × 256 batches. The architecture of the nnU-Net^36^ we used started with an initial layer of two consecutive 3×3×3 convolutions with 32 feature channels, followed by leaky ReLU activation and 2 x 2 x 2 max pooling operations. With each subsequent layer, the number of feature channels was doubled until it reached 320, and the final pooling layer had a kernel size of 1 × 1 × 2. The decoder part of the network utilized 3D-transposed convolution operations to gradually restore the spatial dimensions of the feature map. The final 3 x 3 x 3 convolution reduced the number of channels to seven target categories. A softmax function was then applied to generate a probability distribution of class labels for each voxel. To ensure the robustness of the model, the data was randomly divided into 5 folds, and each fold was trained for 1000 epochs. After training, we ensembled the 5 models. Cross-validation on flies of the same genotype resulted in an average Dice similarity coefficient (DSC) of 0.57 for the 7 classes, with the highest DSC of 0.85 observed for the proventriculus. For the plots in Figure 3, we removed segmentation results which met any one of the following criteria: (1) the volume of any tissue was 0 mm^3^; (2) the volume of any tissue was not a number (NAN); (3) any tissue was segmented into more than 1 class; or (4) any tissue mask was non-contiguous. This filtering resulted in 422 high-quality masks (73% of analyzed 579). The same filtering method was used for Figure 4 in the toxicology screening (see Supplementary Table 1 for filtering results). For the t-SNE plot in Figure 3, we plotted a 3 × 3 inertia tensor. Additional details of the 3D U-Net segmentation algorithm can be found in Supplementary Figure 7 and Supplementary Method 1.

### Random Forest classifier

Downscaled whole-body tomograms were preprocessed into binary segmentations of the wing discs of each animal in order to eliminate the effect of fluorescence intensity differences and emphasize tissue morphology differences. In this preprocessing, we first estimated the bounding box of the larva body. Next, we segmented the body from its surroundings using a Gaussian Mixture Model (GMM). In the final preprocessing step, we localized the wing disc region and segmented it via a *k*-means algorithm. This was followed by a series of postprocessing steps to improve the binary segmentations. From the final binary segmentation of the wing discs, we extracted 30 coordinate-system-free features per image. We analyzed 1,907 tomograms (control *n* = 979; tumorigenic *n* = 928) obtained across 6 dates of measurement and removed 120 resulting failed masks (all voxel values 0) to obtain the 1,787 masks used for downstream analysis. To optimize the Random Forest classifier and evaluate its generalization performance, we employed a nested cross validation with a *k*-replicate strategy to address potential experimental dependencies associated with the measurement date or larva batch. In the nested cross-validation framework, the outer loop was used for assessing the model’s performance, while the inner loop focused on hyperparameter optimization. These tuned hyperparameters included the maximum number of splits in the tree-based model, the number of trees, and the number of features to be selected at each split. Specifically, the maximum number of splits was varied across the set {3,5,8,10,20,30,40,50,100}, the number of trees across the set {5,10,20,30,40,50,100,200,300,400,500,1000}, and the number of selected features per split across the set {4, 5, 6, 7, 8,9,10,20}. Within the internal cross validation loop, each validation cycle involved selecting five out the six measurement dates. Among these selected dates, four were allocated for the training set, and one was designed for the validation where the model was predicted. Consequently, for each set of selected hyperparameters, the model underwent training and validation five times with different combinations of measurement dates. The optimal set of hyperparameters was identified by maximizing the average area under the ROC curve (AUC) obtained over the 5 validations set. The set of the best hyperparameters was then used to build the model by training on the entire training set comprised of 5 measurement dates and tested on the external test set. This entire procedure was iteratively conducted six times, corresponding to the total number of measurement dates, ensuring that each date was used once as the external test set. The performance of the Random Forest classifier on the test set was as follows: average accuracy 67.85% with a standard deviation (+/-) of 5.42%; average F1 score 66.05% (+/- 9.36%); average recall of 65.72% (+/- 8.21%); average precision of 68.18% (+/- 18.76%); and average AUC of 76.10% (+/- 5.52%). The detailed protocol including preprocessing, segmentation, feature extraction, and classification can be found in Supplementary Method 2. While imaging for classifier training, we flowed and imaged 1,041 flies in 23 hr.

### Fly food for toxicological screening

To prepare control food, agar (0.375 g, Wako), glucose (4 g, Wako), cornmeal (2 g, OYC Americas, Inc.), dry yeast (4 g, Asahi Group Foods, Ltd.), propionic acid (0.25 mL, Wako), and 70% ethanol (0.25 mL, Wako) were added to Milli-Q purified water. To prepare food infused with 500 nm and 4 µm PS particles: 6.34 mL of a 2.5 wt% PS beads suspension (Wuxi Rigor Technology Co, Ltd.) was added to the control food mixture to make the 2.6 µg/g food; and 6.34 mL of a 10× diluted 2.5 wt% PS beads suspension was added to the control food mixture to make the 0.26 µg/g food. To prepare food infused with 15 µm PMMA particles: 152.3 mg of powder PMMA particles (Japan Exlan Company, Ltd.) was added to the control food mixture to make the 2.5 µg/g mM food; 15.23 mg of powder PMMA particles (Japan Exlan Company, Ltd.) was added to the control food mixture to make the 0.25 µg/g food. To prepare food infused with 6 µm PTFE particles: 152.18 mg of powder PTFE particles (AS ONE Co.) was added to the control food mixture to make the 2.5 µg/g food; 15.218 mg of powder PMMA particles (AS ONE Co.) was added to the control food mixture to make the 0.25 µg/g food. Each food mixture was boiled and cooled down after mixing the ingredients.

### Statistical analyses and toxicological screening

The statistical analyses of results from the *nub-GAL4* basic performance demonstration (Figure 3) and MP toxicology experiment (Figure 4) were performed using Python 3. Measurements were taken from distinct samples. We performed an unpaired t test (two-sided), with *m* + *n* - 2 degrees of freedom, where *m* and *n* are the sample sizes of the two comparison groups. A *p* value of over 0.05 is designated as “ns”; *p* < 0.05 is “*”; *p* < 0.01 is “**”; and *p* < 0.001 is “***” (see Supplementary Table 5 for Cohen’s *d* calculations). For the MP experiment, all samples were the same strain of *nub-GAL4* flies and were cultured in the same incubator at the same temperature (24 °C). Each condition was comprised of larvae from 2 separate food tubes. Flipping of flies was performed every 2 days. Larvae from each cohort (control, 0.25 or 0.26 µg/g, 2.5 or 2.6 µg/g) were collected at the same time. Fixation and tissue clearing were performed at the same time, such that all flies in a cohort had common exposure to the chemicals. Imaging of each cohort was performed under the same conditions of laser power, LSM alignment, room temperature, and room humidity.

### Genotypic screening

For genotypic screening, the mixed-genotype progenies were manually separated into target and non-target cohorts. Samples from each pure cohort were loaded into the flow zoometer for calibration rapid image acquisitions to determine the sorting criteria. Following the calibration, samples from the two cohorts were manually mixed in equal portions and loaded into the flow zoometer. In order to differentiate between target and non-target flies, each whole larva was rapidly scanned (at a rate of 2 mm/sec or 100 µm/frame) by the flow zoometer. In body-length-based screening, the rapid scan was performed with the blue channel, and the body length was estimated from the raw image stack by counting the frames whose mean intensity surpassed a threshold value set near the ambient light background level. The system compared the estimated body length from each rapid scan to the threshold value determined in the calibration experiment and identified the sample as a target or non-target. The total screening time for each larva was 10.5 sec (7 sec for the rapid scan, 3.5 sec for decision making). In fluorescence-based genotypic screening, the rapid scan was performed in the green channel. The presence of the mCherry expression was evaluated by comparison of the image stack frames to a threshold value determined in the calibration experiment, and the sample was identified as a target or non-target. The total screening time for each larva was 10.9 sec (7 sec for the rapid scan, 3.9 sec for decision making). Target samples were subsequently imaged at a high axial resolution (scan speed 0.2 mm/sec or 10 µm/frame), while non-target samples were released without imaging again. The purity was calculated as the number of high-resolution images of target genotypes divided by the total number of high-resolution images. The yield was calculated as the number of high-resolution images of target genotypes divided by the total number of target genotypes loaded into the flow zoometer.

### Fly stocks

The following *Drosophila* strains were obtained from the Bloomington *Drosophila* Stock Center: *nub-GAL4* (#86108), *nub-GAL4,UAS-myr::mRFP* (#63148), *UAS-2xEGFP* (#6874), *UAS-mCherry.NLS* (#38424), *UAS-mtdTomato,10xQUAS-6xGFP^3xP3-RFP^/CyO;r4-QF2/TM6B* (#66476), *GMR-GAL4/CyO* (#9146), *GMR-GAL4/CyO-Tb-RFP* (created in this study), *UAS-mCherry.CAAX* (#59021); *Drosophila* Genomics and Genetic Resources: *w^1118^* (#123720). Additional strains used were: *ppk-GAL4*,*P{UAS-mCD8::GFP.L}Ptp4E^LL4^* on X chromosome^35^, *Ubx-GAL4* (from Dr. L. S. Shashidhara, Ashoka University, India), *Ubx-GAL4/TM6B* (created in this study), *UAS-Ras*^G12D^,*UAS-p53*^shRNA^,*UAS-Med*^shRNA^,*UAS-dCycE* (*UAS-4-hit*)^40,41^, *ptc-GAL4,UAS-mCherry;+/SM5_tubP-GAL80_-TM6B* (created in this study).

### Fly food for maintaining stocks

Brewers’ yeast (MP Biomedicals), yeast extract (Sigma Aldrich), bacto casitone (BD), agar, sucrose, glucose, MgCl_2_, CaCl_2_, propionic acid and mold inhibitor (10% methyl-4-hydroxybenzoate in 95% ethanol) (Fujifilm Wako) were dissolved in water purified with reverse osmosis filters. The mixture was then boiled and cooled down.

### Confocal microscopy of wing discs

Larval wing discs were dissected out of L3 larvae and fixed in 4% PFA in PBS for 20 min. After washing in PBT, the discs were incubated overnight in DAPI solution (Fujifilm Wako) diluted with PBS (-) at 1:1000. Then the discs were mounted in Vectashield Vibrance Antifade Mounting Medium (Vector Biolabs) and imaged with a Leica TCS SPE confocal microscope (Leica Microsystems).

### Flies for genotypic screening

*GMR-GAL4* balanced with *CyO-Tb-RFP* were crossed with *UAS-mCherry.CAAX* at 25° C to produce flies for fluorescence-based screening: *GMR-GAL4/UAS-mCherry.CAAX* (target; mCherry expression in eye discs) and *+/CyO-Tb-RFP* (non-target; no mCherry expression). *Ubx-GAL4* balanced with *TM6B* were crossed with *UAS-mCherry.NLS* at 25° C to produce flies for body length-based screening: *Ubx-GAL4/UAS-mCherry.NLS* (target; non-tubby phenotype) and *UAS-mCherry.NLS/TM6B* (non-target; tubby phenotype).

### Flies for tumor screening

Virgin females of *ptc-GAL4,UAS-mCherry;UAS-4-hit/SM5_tubP-GAL80_-TM6B* were crossed with *w^-^* males to obtain *ptc-GAL4,UAS-mCherry/+*;*UAS-4-hit/+* (*ptc*>*4-hit*) offspring. Resulting progenies were cultured until L3 larvae at 22 °C or 23 °C.

## Acknowledgements

This work was supported by the JST START Program (JPMJST2115), JST CREST Program (JPMJCR2333), AMED Drug Discovery Platform Promotion Research Program (22ak0101163h0002), AMED Project for Promotion of Cancer Research and Therapeutic Evolution (23ama221229h0001), JSPS Core-to-Core Program (JPJSCCA20190007), JSPS KAKENHI (19H05412, 20H03524), Whiterock Foundation, G-7 Scholarship Foundation, and the Junior Scientist Promotion and Photo-Excitonix at Hokkaido University. Our sincere appreciation goes to T. Ota, who assisted in creating the digital switches for the DC power supply in the flow zoometer pinch valves and peristaltic pumps. We are grateful to M. Sehara for her contributions in manual tissue clearing. We thank H. Terakawa for providing mock-up illustrations for the 3D animated video and Didan and his team for animating the 3D video. We extend our gratitude to S. Mori and M. Yamada for their technical support in *Drosophila* husbandry.

## Contributions

K.G. conceived of the concept of flow zoometry. K.G., M.S., and T.Kom. conceived of the project. W.P., Z.H., and K.Hir. designed the automated tissue-clearing device. W.P. and Z.H. built the automated tissue clearing device. W.P. and H.W. programmed the robotic arm. T.Kob., W.P., Z.H., Y.N., S.A., H.W., K.Hir., and L.K. designed the larva transport and flow system. T.Kob., W.P., S.A., and L.K. built the larva transport and flow system. K.G., K.Hir., L.K., P.M., H.K., K.Hua., and W.P. designed the optical 3D microscope. K.Hir., L.K., P.M., and W.P. built the 3D microscope. W.P. and H.W. designed and built the software for the large-volume LSM. W.P. built the software for data processing and management. K.Hir., W.P., M.Her., and J.A. designed and wrote the code for tomographic image reconstruction. C.Z. designed, built, and trained the 3D U-Net segmentation algorithm. W.P., Y.Nag., and C.H. produced manual annotations of 3D tomograms for the 3D U-Net segmentation algorithm training. J-E.C., J.A., and T.Kom. designed, built, and trained the segmentation and classifier algorithms for tumor screening. H.W., C.H., W.P., and T.Kob. carried out imaging to demonstrate the basic performance of flow zoometry. H.W., W.P., T.Kob., and H.K., carried out imaging for training the 3D U-Net segmentation algorithm. T.Kob. designed and realized the MP toxicological screening including sample handling, tissue clearing, imaging, and analysis. H.W., S.Y., and W.P. designed genotypic screening. H.W. carried out programming, imaging, and analysis for genotypic screening. M.S., S.H., T.O., Y.W., W.P., K.G., J.A., J-E.C., and R.Y. designed the tumor screening. S.H., T.O., Y.W., S.M., R.Y., H.U., E.K., and M.S. directed the fly genetic design and breeding schemes. S.H., T.O., Y.W., S.M., R.Y., T.Kob., H.K., W.P., H.U., K.F., and H.W. carried out fly crossbreeding and sample preparation. H.W., T.K., C.H., W.P., L.K., P.M., and K.Hir. carried out other imaging experiments. All authors contributed to interpreting experimental results. W.P., M.S., J-E.C., O.K., S.Y., H.B., C.Z., K.I., T.I., H.U., E.K., Y.W., T.Kom., and K.G. wrote the manuscript. W.P., T.Kob., C.Z., J-E.C., Z.H., H.W., J.A., M.Hay., Y.Nak., and K.G. composed the figures. W.P., K.Hir., M.S., T.Kom., and K.G. led the project. K.G., M.S., and T.Kom acquired funding for the project.

## Ethics statement for the use of genetically modified organisms

Approval for the use of genetically modified organisms was provided by the Japanese Ministry of the Environment, via the University of Tokyo Expert Committee on Genetically Modified Organisms and the Hokkaido University Safety Committee on Genetic Recombination Experiments (approval numbers: 2019–007 and 2022–029), in accordance with the Act on the Conservation and Sustainable Use of Biological Diversity through Regulations on the Use of Living Modified Organisms (Act No. 97 of 2003) and The Ministerial Ordinance Providing Containment Measures to Be Taken in Type 2 Use of Living Modified Organisms for Research and Development.

---

**Supplementary Video 1.**
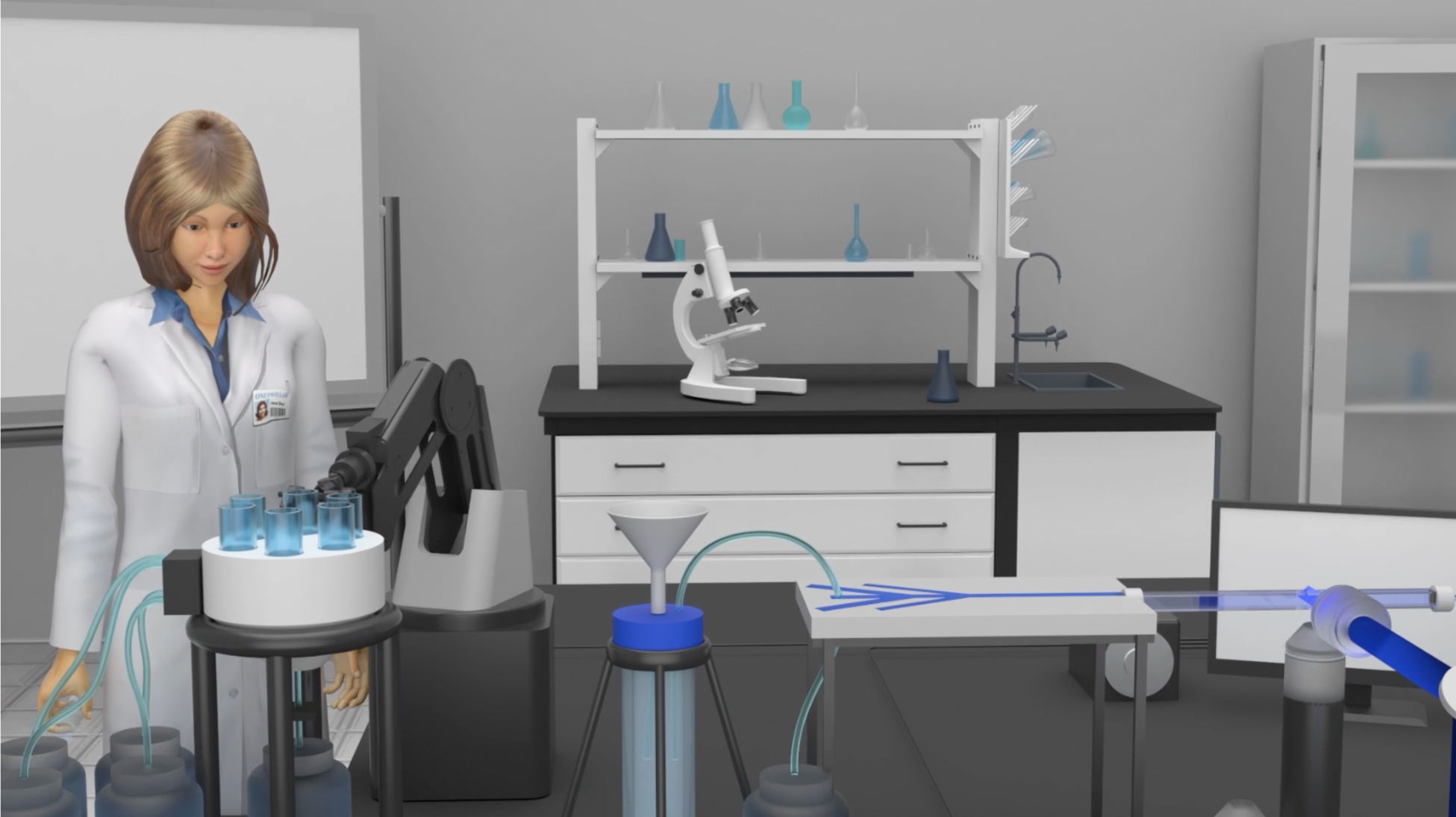
Animated video of flow zoometry. The flow zoometer performs fixation, tissue clearing, transport and isolation, tomographic imaging, and data-driven analysis of *Drosophila* L3 larvae in a seamless, fully automated manner. The developed system can tissue-clear over 1,000 larvae within 24 hr and subsequently obtain over 1,000 3D whole-body tomograms at single-cell resolution within 24 hr. The music was composed and recorded by W.P.

**Supplementary Video 2.**
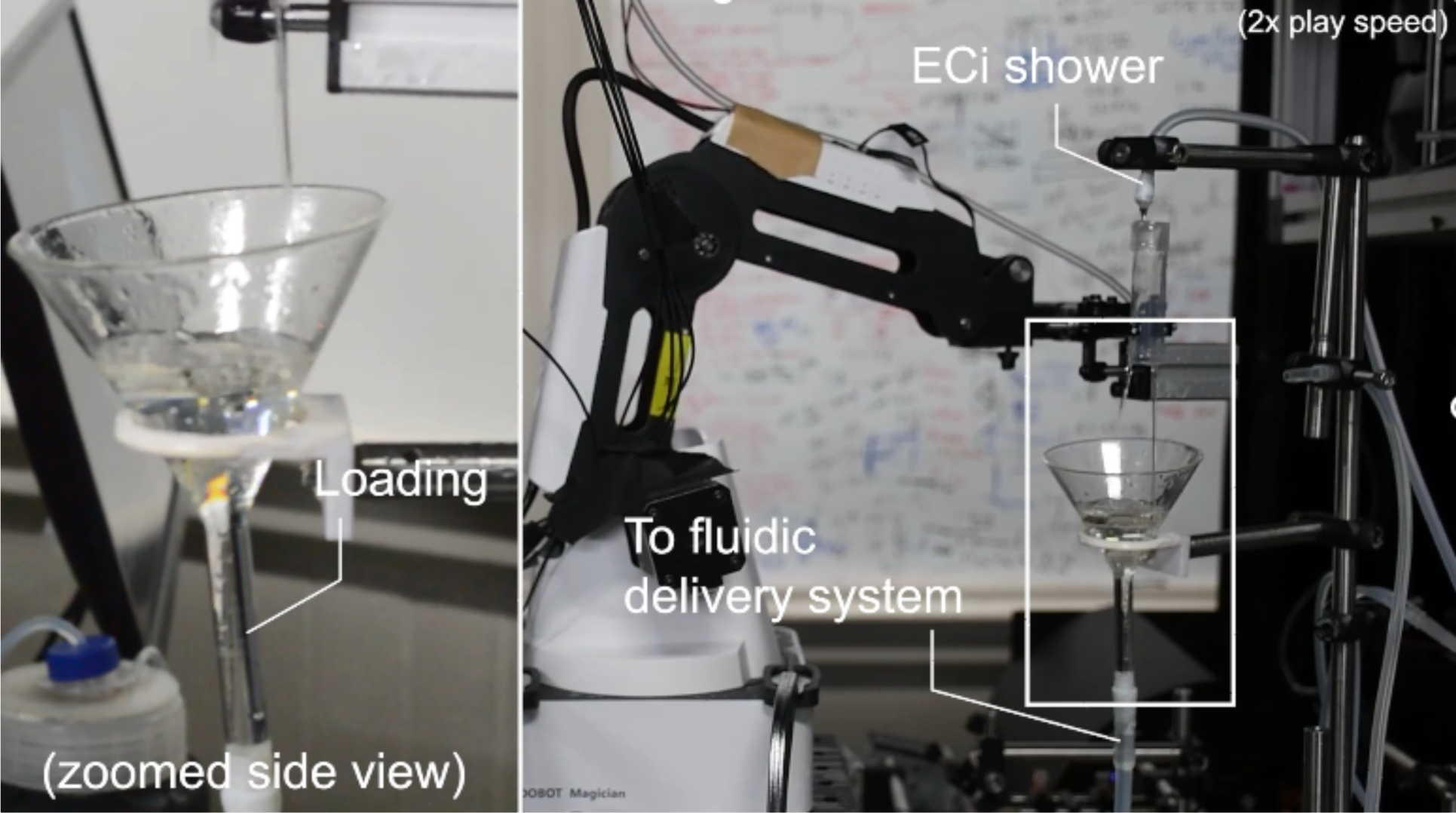
Operation of the flow zoometer. Scenes of a flow zoometry experiment show the operation of the tissue-clearing device, the robotic sample loader, the fluidic delivery system, and the large-volume LSM in sequence.

**Supplementary Video 3.**
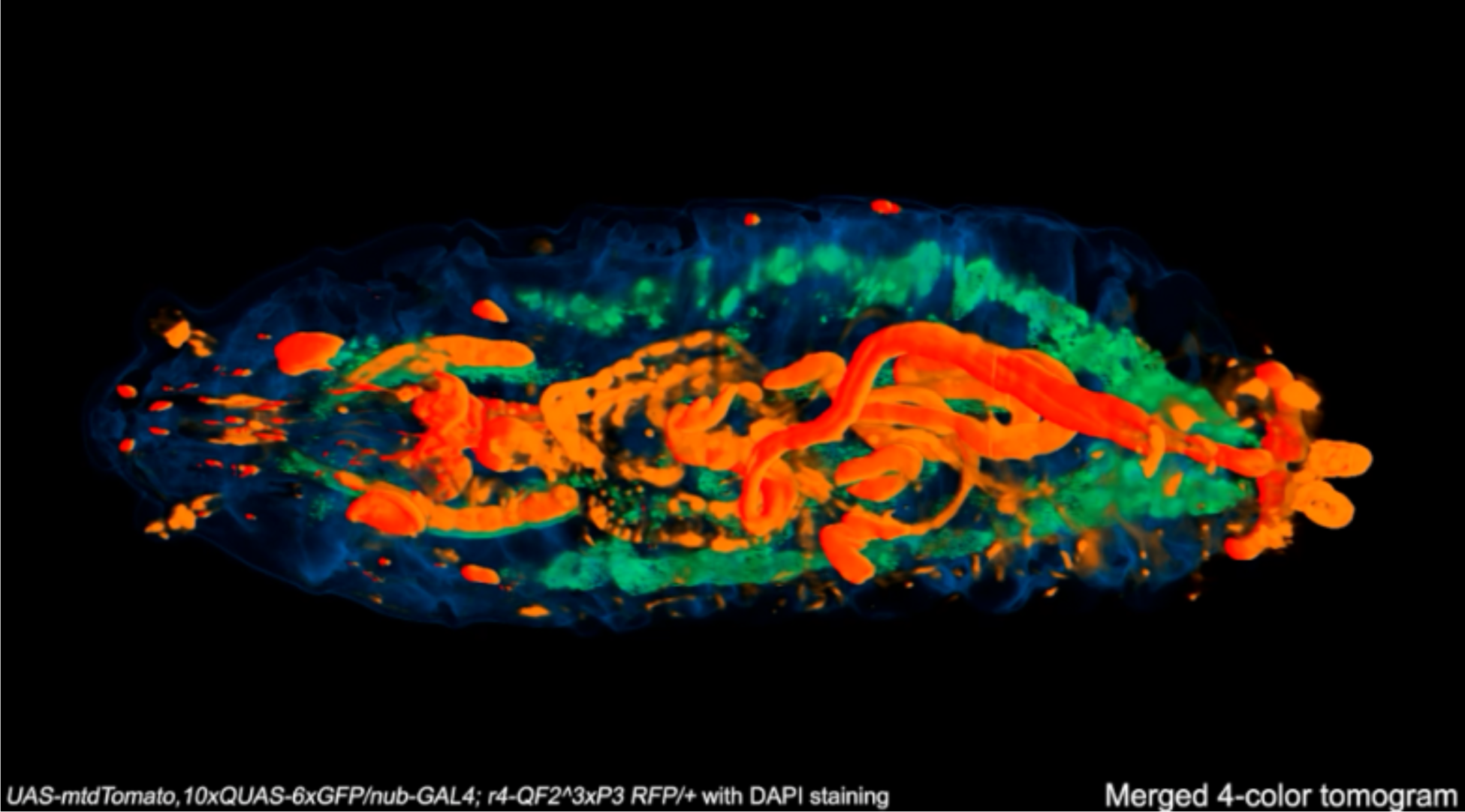
Whole-body 3D tomograms of *Drosophila* acquired by the flow zoometer. Shown in sequence are several animations of tomograms of *Drosophila* larva obtained by flow zoometry. In the whole-body 3D image data, individual internal tissues expressing fluorescent proteins are resolved with high contrast against background autofluorescence.

**Supplementary Video 4.**
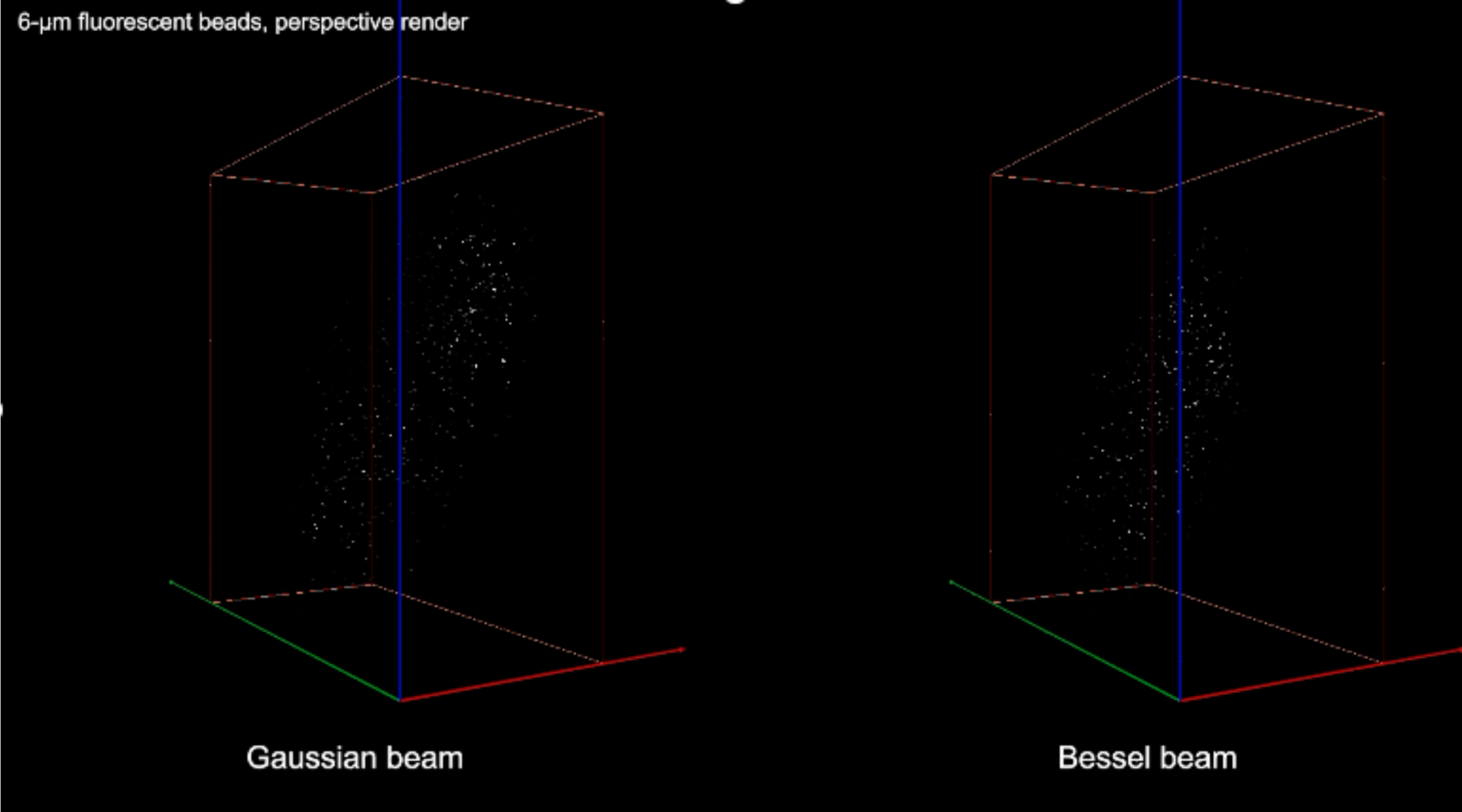
Tomograms of 750 nm and 6 µm fluorescent microspheres acquired by the flow zoometer. Shown in the video are tomograms of green-fluorescent 750 nm and 6 µm PS microspheres suspended in glycerol obtained with the Gaussian and Bessel beams by the flow zoometer. First, perspective renderings of the tomograms (Amira 2022.2 volren, linear greyscale) are shown rotating in space about 2 axes. The bounding boxes are displayed in orange, and the *x*, *y*, and *z* axes are displayed in green, blue, and red, respectively. The bounding box volume is 2.42 mm × 2.80 mm × 1.21 mm for all tomograms. Second, orthographic renderings (*xy* plane) are shown at different zoom levels with scale bars.

**Supplementary Figure 1.**
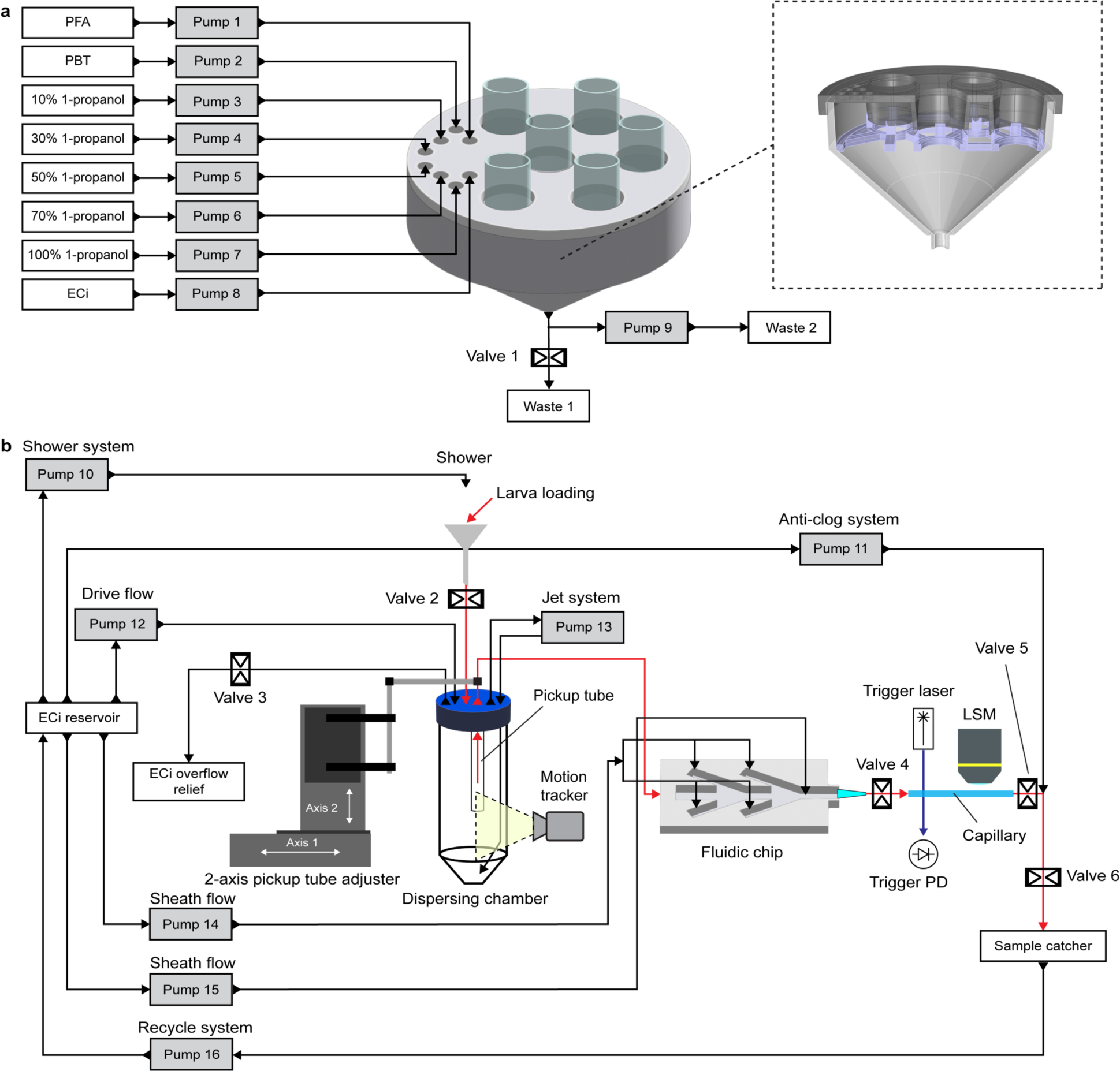
Details of the tissue-clearing device and fluidic delivery system. **a,** Detailed schematic of the tissue-clearing device. Reagents for tissue clearing are delivered in sequence to samples via an array of 8 peristaltic pumps (grey boxes, Pumps 1 to 8). Used reagents are evacuated from the main chamber via an outlet at its bottom. The used reagents can be sorted (Valve 1, Pump 9) according to their type to multiple waste tanks. In the dashed box to the right is a cross sectional view of the main chamber which shows its construction. The main chamber comprises 3 3D-printed parts and is designed to firmly hold sample tubes in place and enable the unhindered delivery of the tissue-clearing reagents to the larvae. The design furthermore facilitates the pickup of sample tubes by the robotic sample loader. **b,** Detailed schematic of the fluidic delivery system. The path of larvae (suspended in ECi) throughout the system is indicated by red arrows, while paths of ECi flowed through the system are indicated by black arrows. Pump systems or functions are noted above the pumps. Larvae are loaded into the system via a funnel and Valve 2 while a shower system (Pump 10) ensures all larvae enter the system, after which Valve 2 is closed and Pump 10 is stopped. In the dispersing chamber, larvae are circulated via a jet system (Pump 13). A drive flow (Pump 12) from the ECi reservoir into the dispersing chamber pushes individual larvae and small larval aggregates out of the dispersing chamber and into the fluidic chip. In the fluidic chip, sheath flows (Pumps 14 and 15) separate larval aggregates while individual larvae are oriented lengthwise with respect to the flow direction. Larvae exit the dispersing chamber and enter the LSM capillary. A 450 nm continuous wave trigger laser and photodetector (PD) are used to detect the entry of larvae into the capillary. When a larva is detected, Valves 4 and 5, respectively before and after the capillary, are closed, and all Pumps are stopped, completely immobilizing the larva in the capillary. After imaging, Valves 4 and 5 are opened and Pumps 12 to 15 are initiated. Used larvae are rescued in a mesh filter-based sample catcher, while ECi is returned to the ECi reservoir via the recycle system (Pump 16). A video camera-based motion tracker is used to detect (i) if larvae are exiting the dispersing chamber and (ii) the presence of remaining samples. This information is used in combination with the frequency of sample detection by trigger laser and PD, to determine (i) if a clog has occurred or (ii) if it is time to load the next sample. If a clog has been detected, Valve 6 closes, Valve 3 opens, and the anti-clog system (Pump 11) is initiated to reverse the system flow and send clogged larvae back to the dispersing chamber. Excess ECi is dumped into and ECi overflow tank.

**Supplementary Figure 2.**
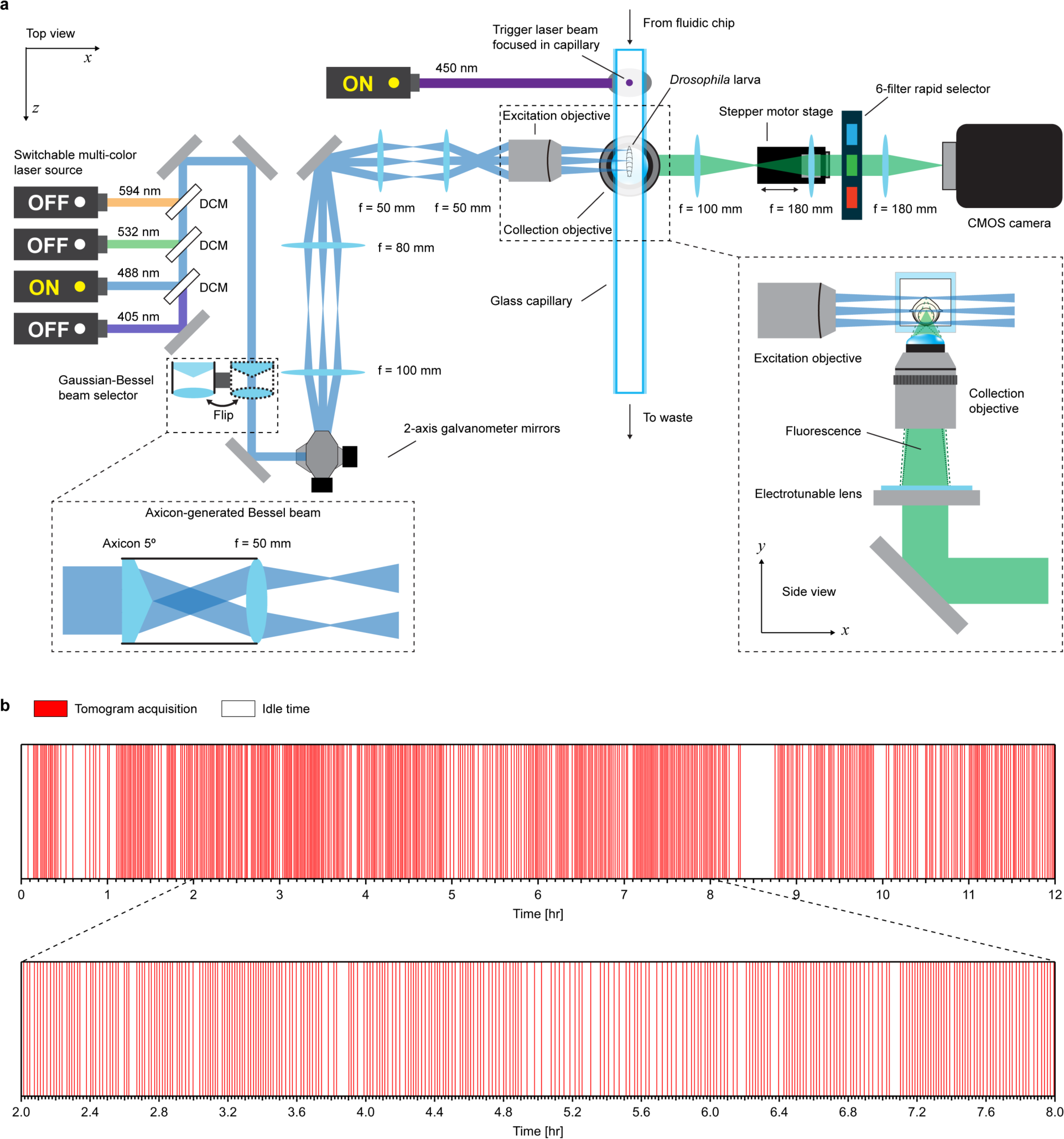
Structure and function of the large-volume LSM. **a,** Schematic of the large-volume LSM. A switchable multi-color laser source generates a Gaussian laser beam which is converted into a scanned light-sheet via a 2-axis galvanometer mirror scanner, relay lenses, and an excitation objective lens. An axicon and lens system can be flipped in the Gaussian beam path to generate a Bessel beam at the sample position. The digital scanned light-sheet is focused onto the interior center of a square-cross-section glass capillary, where *Drosophila* L3 larvae are immobilized in ECi after their presence is sensed by a trigger laser beam focused in the capillary upstream. The capillary is mechanically scanned through the immobilized larva for each color. Fluorescence is collected via a collection objective lens, electrotunable lens, 6-filter rapid selector, and CMOS camera. A lens mounted on a stepper motor stage is used to adjust the focus of the fluorescence on the CMOS camera. After imaging, the larva is release to the waste. DCM, dichroic mirror. **b,** Time-series representation of 3D tomogram acquisitions illustrating the throughput of the large-volume LSM. Image acquisitions are shown in red, while idle time (e.g., flowing of larvae through the LSM) is shown in white. Using the robotic sample loader for automation, we obtained 505 3D tomograms within a span of 12 hr. Idle times vary due to both the stochastic nature of sample picking in the dispersing chamber as well as clogging and the subsequent engagement of the automated anticlogging system. The image acquisition time was defined as the raw tomogram data file creation timestamp, and the plot is a histogram of all timestamps. The bin width is approximately 5 sec, such that all timestamps are represented by individual lines in the plot.

**Supplementary Figure 3.**
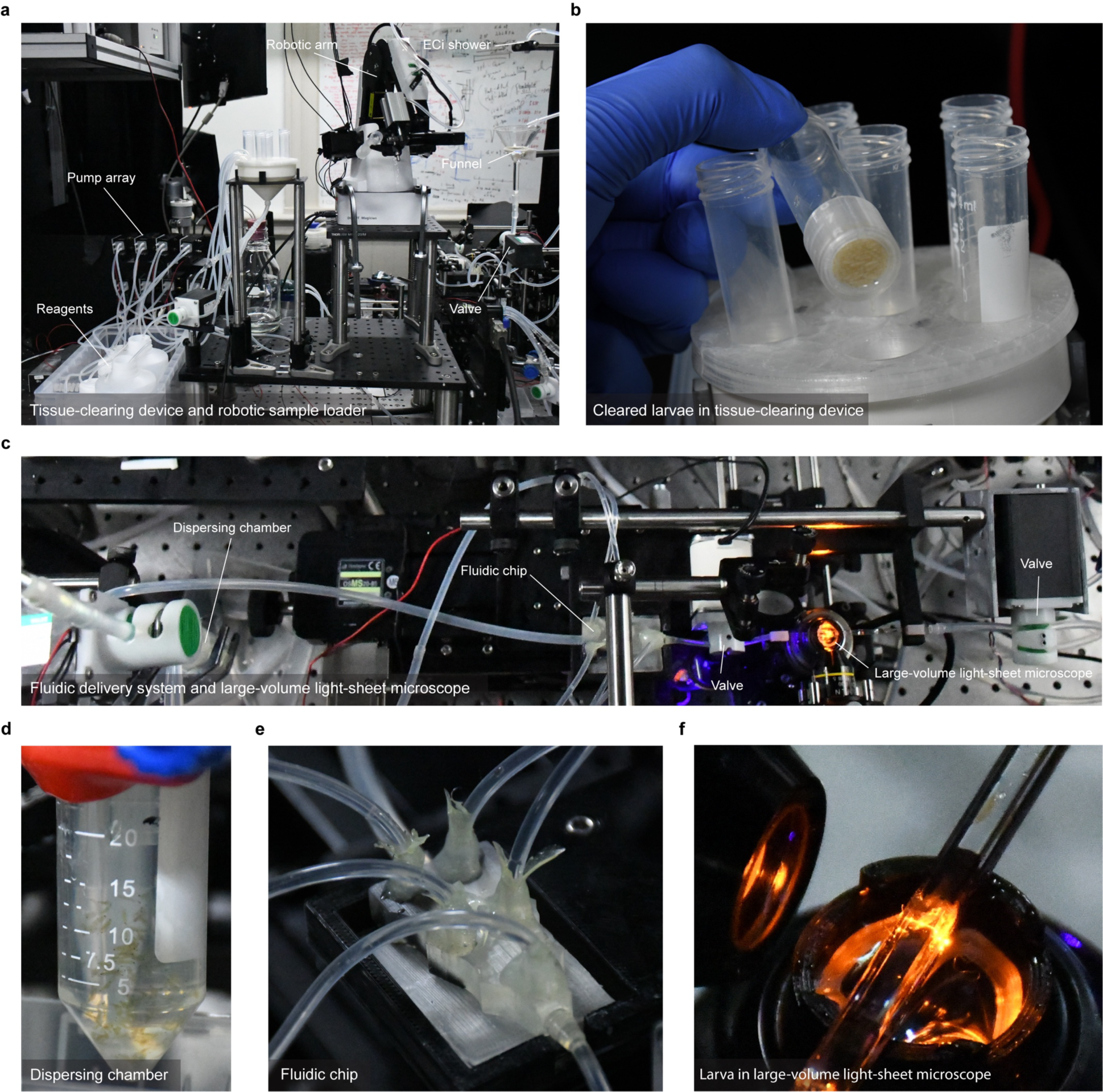
Pictures of the flow zoometer. **a,** Pictures of the tissue-clearing device (left side of picture) and robotic sample loader (right side of picture). In the left foreground are the reagent stocks and pump array of the tissue-clearing device, while the main chamber of the tissue-clearing device is near the center foreground. To the right of the tissue-clearing device is the robotic arm of the robotic sample loader. In the far right side of the picture is the ECi shower, funnel, and valve of the robotic sample loader, which leads to the fluidic delivery system. **b,** Close-up picture of the tissue-clearing device showing cleared larvae in one of the sample tubes. **c,** Picture of the fluidic delivery system and large-volume LSM. Flow proceeds from the dispersing chamber on the left toward large-volume LSM on the right. The 2 pinch valves flanking the capillary in the large-volume LSM ensure that larvae are immobilized during image acquisition. **d,** Close-up picture of the dispersing chamber with larvae in circulation. **e,** Close-up picture of the fluidic chip. Larvae enter the chip via the large tube in the background (coplanar with the chip), while the 5 narrower tubes (non-coplanar with the chip) provide sheath flows which isolate and align individual larvae during their passage through the fluidic chip. **f,** Close-up picture of a larva in the large-volume LSM. The custom-made black ring beneath the capillary ensures that the immersion liquid of the collection objective is retained during long flow zoometry experiments.

**Supplementary Figure 4.**
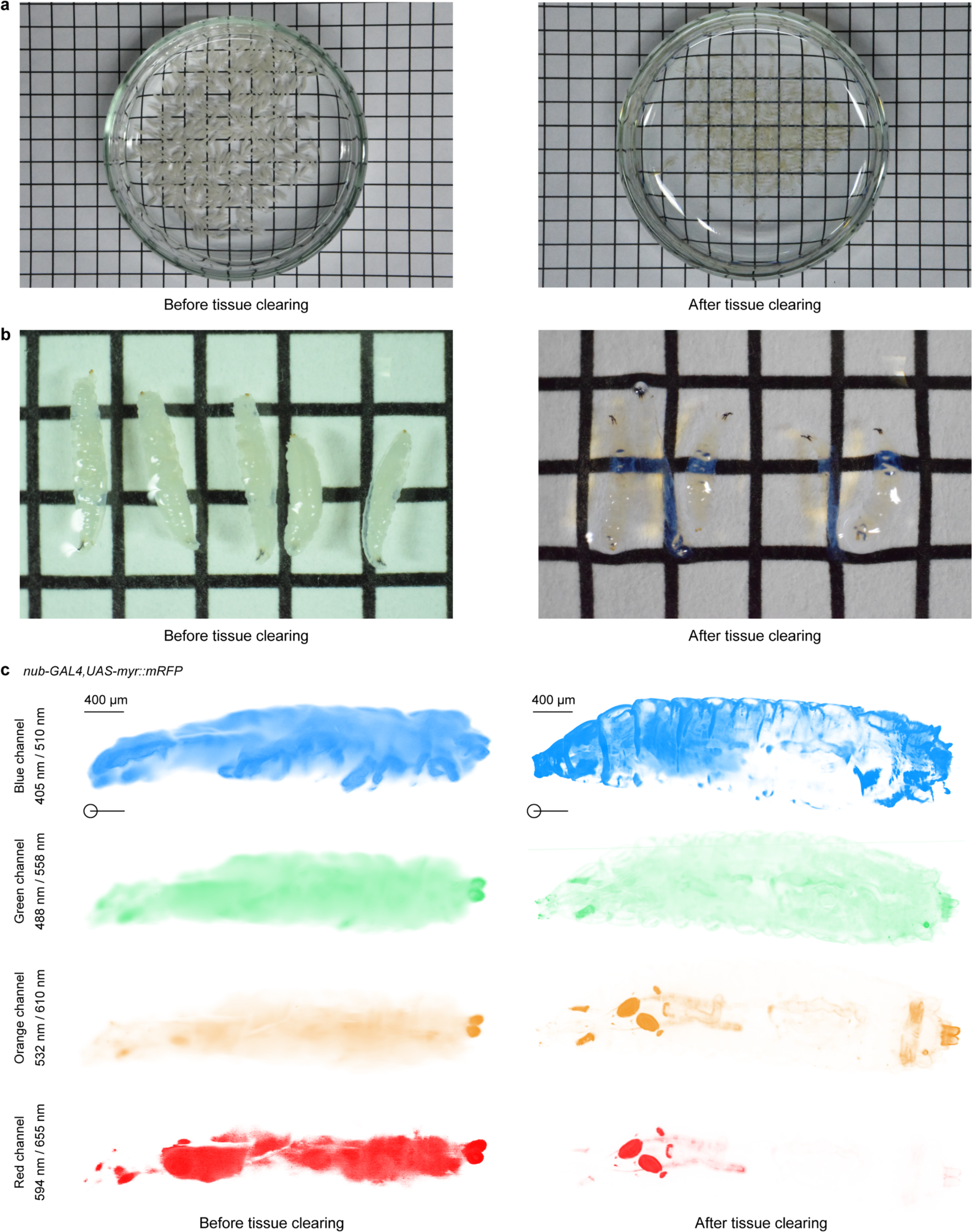
Before and after tissue clearing of *Drosophila* larvae. **a,** Pictures of >100 fixed, uncleared *Drosophila* L3 larvae before dehydration and refractive index matching (left) and >100 cleared *Drosophila* L3 larvae after dehydration and refractive index matching (right). Grid spacing 0.5 cm. **b,** Microscopy images of fixed, uncleared *Drosophila* L3 larvae before dehydration and refractive index matching (left) and cleared *Drosophila* L3 larvae after dehydration and refractive index matching (right). Grid spacing: 0.25 cm. **c,** Representative 4-color tomograms of a fixed, uncleared *Drosophila* L3 larva before dehydration and refractive index matching (left) and a cleared *Drosophila* L3 larva after dehydration and refractive index matching (right). Before tissue clearing, RFP-expressing tissues such as the wing discs are not detected. After tissue clearing, RFP-expressing tissues are detected with high contrast. Tomograms are *xz* orthographic projections displayed in Amira’s Volren mode, with complete opacity. Intensity ranges are displayed from 180 (to remove the ambient light background) to 1/4 of the maximum intensity.

**Supplementary Figure 5.**
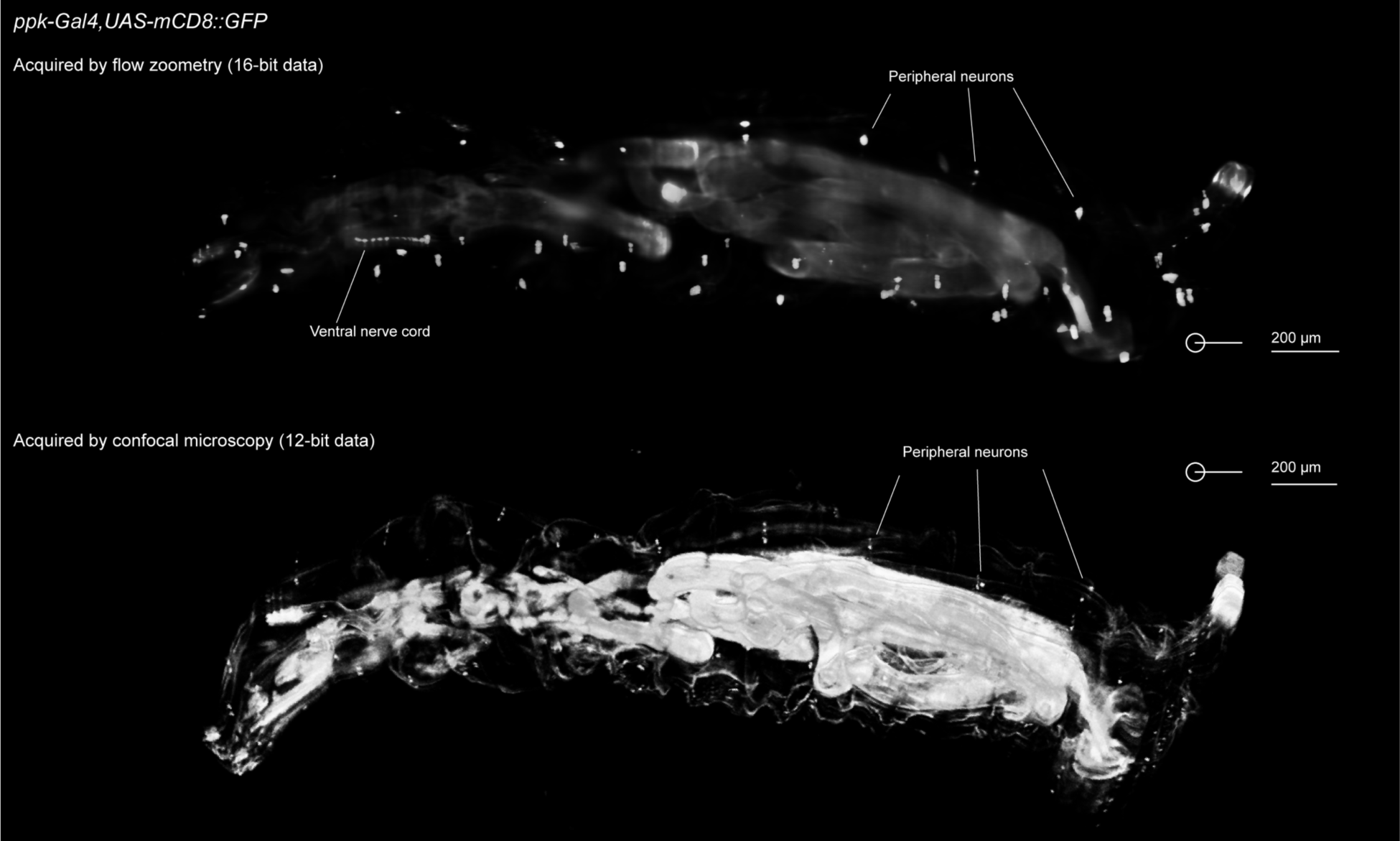
Comparison of 3D tomograms of the same *Drosophila* larvae obtained by the flow zoometer and by confocal fluorescence microscopy. Shown is a 3D tomogram of the same *ppk-GAL4,UAS-mCD8::GFP* larva obtained by the flow zoometer (upper) and by a confocal fluorescence microscope (lower). The comparison demonstrates that flow zoometry accurately images the whole larval body, with respect to confocal microscopy. The pattern of peripheral neurons on the larval body is overall conserved between the two images. The perspective of both images is orthogonal to the lateral plane (*xz*). Some differences in the displayed fluorescence intensities between the two images can be attributed to the data bit depth, in addition to the imaging equipment. The scale bars were calculated independently using 6 µm fluorescent beads for scale calibration.

**Supplementary Figure 6.**
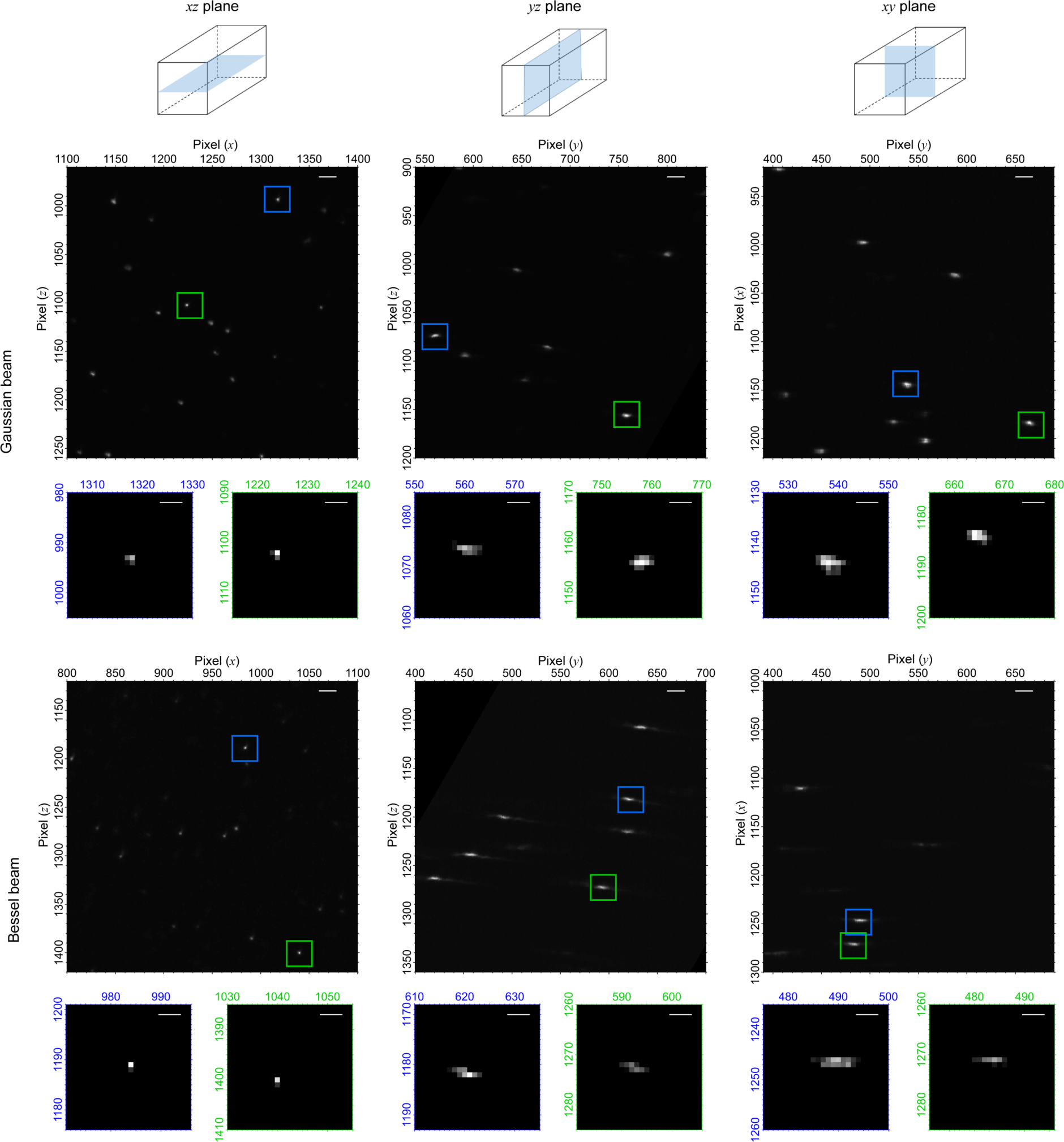
Spatial resolution of the flow zoometer. Obtained 3D tomograms of 750 nm fluorescent PS microspheres suspended in glycerol are shown. Pictographs at the top indicate the displayed plane, *xy*, *yz*, and *xz* from left to right. Results are shown for both Gaussian (upper row of 3 sets of 3 plots) and Bessel (lower row of 3 sets of 3 plots) beams in the large-volume LSM. For each beam, below each larger plot are 2 insets of zoomed-in regions corresponding to the colored boxes in the larger plot. In the 3 larger plots for each beam type, the full intensity range is displayed. In the insets, the plotted intensity ranges from the half-maximum value to the maximum value. In all plots, the pixel side lengths were 1.1 µm and the scanning speed was 1 µm/frame. Larger plot scale bars: 20 µm; inset scale bars: 5 µm.

**Supplementary Figure 7.**
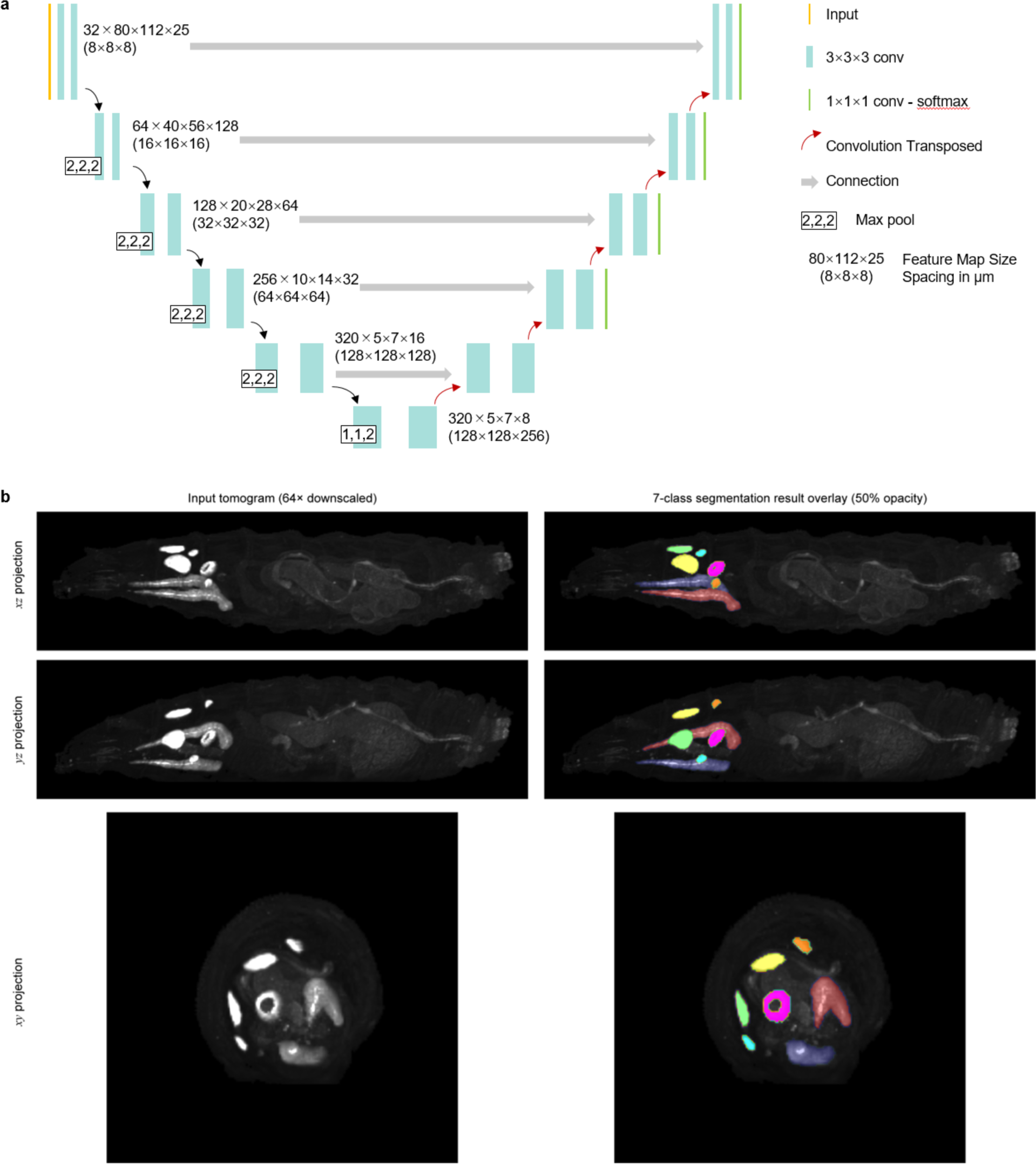
Structure and performance of the 3D U-Net segmentation algorithm. **a,** Structure of the 3D U-Net segmentation algorithm. **b,** Multi-axis orthographic projections of a representative input larva tomogram (*nub-GAL4,UAS-myr::mRFP*; left) and resulting 7-class segmentation mask (shown with 50% opacity overlaid on the input tomogram; right) calculated by the 3D U-Net segmentation algorithm. All 7 classes of tissues of interest including the left wing disc, right wing disc, left haltere disc, right haltere disc, left salivary gland, right salivary gland, and proventriculus are identified clearly by the algorithm.

**Supplementary Figure 8.**
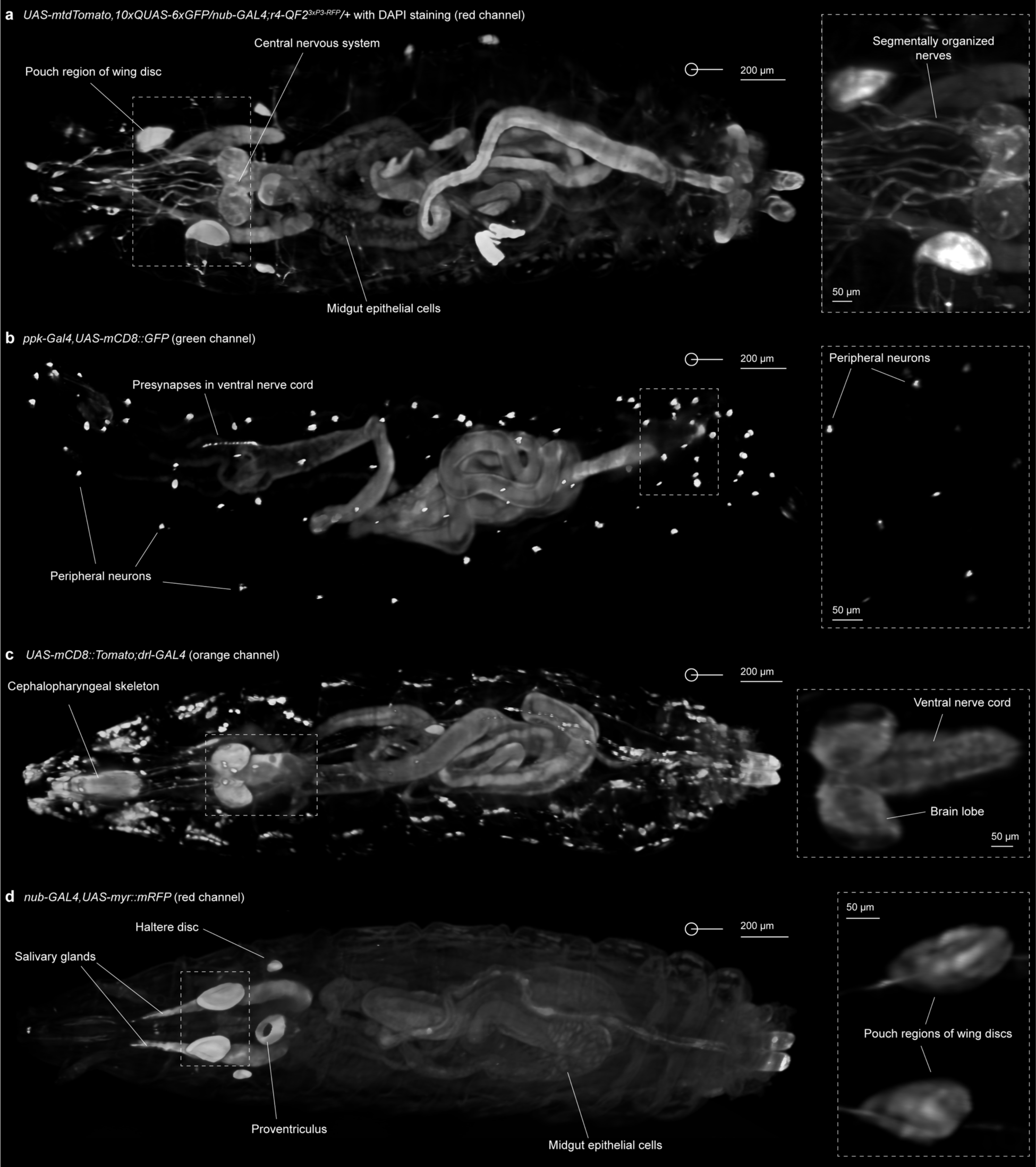
3D tomograms in greyscale of various types of *Drosophila*. **a,** Tomogram (red channel) of a 4-color larva corresponding to that shown in Figure 2a. RFP expression highlights multiple tissues including segmentally organized nerves, the central nervous system, the pouch regions of the wing discs, and the midgut epithelial cells. **b,** Tomogram (green channel) of a *ppk-GAL4* larva corresponding to that shown in Figure 2b. GFP expression occurs in peripheral neurons and presynapses in the ventral nerve cord. **c,** Tomogram (orange channel) of a *drl-GAL4* larva corresponding to that shown in Figure 2c. TdTomato is expressed in the cephalopharyngeal skeleton, the central nervous system including the brain and ventral nerve cord, and the segmentally organized nerves. **d,** Tomogram (red channel) of *nub-GAL4* larva corresponding to that shown in Figure 2d. The expression of mRFP is evident in the haltere discs, pouch regions of the wing discs, salivary glands, proventriculus, midgut epithelial cells, and anal plate. All images are orthographic *xz* projections with voxel opacity set to 1. The intensity ranges of the insets were adjusted to highlight finely resolved tissue features.

**Supplementary Figure 9.**
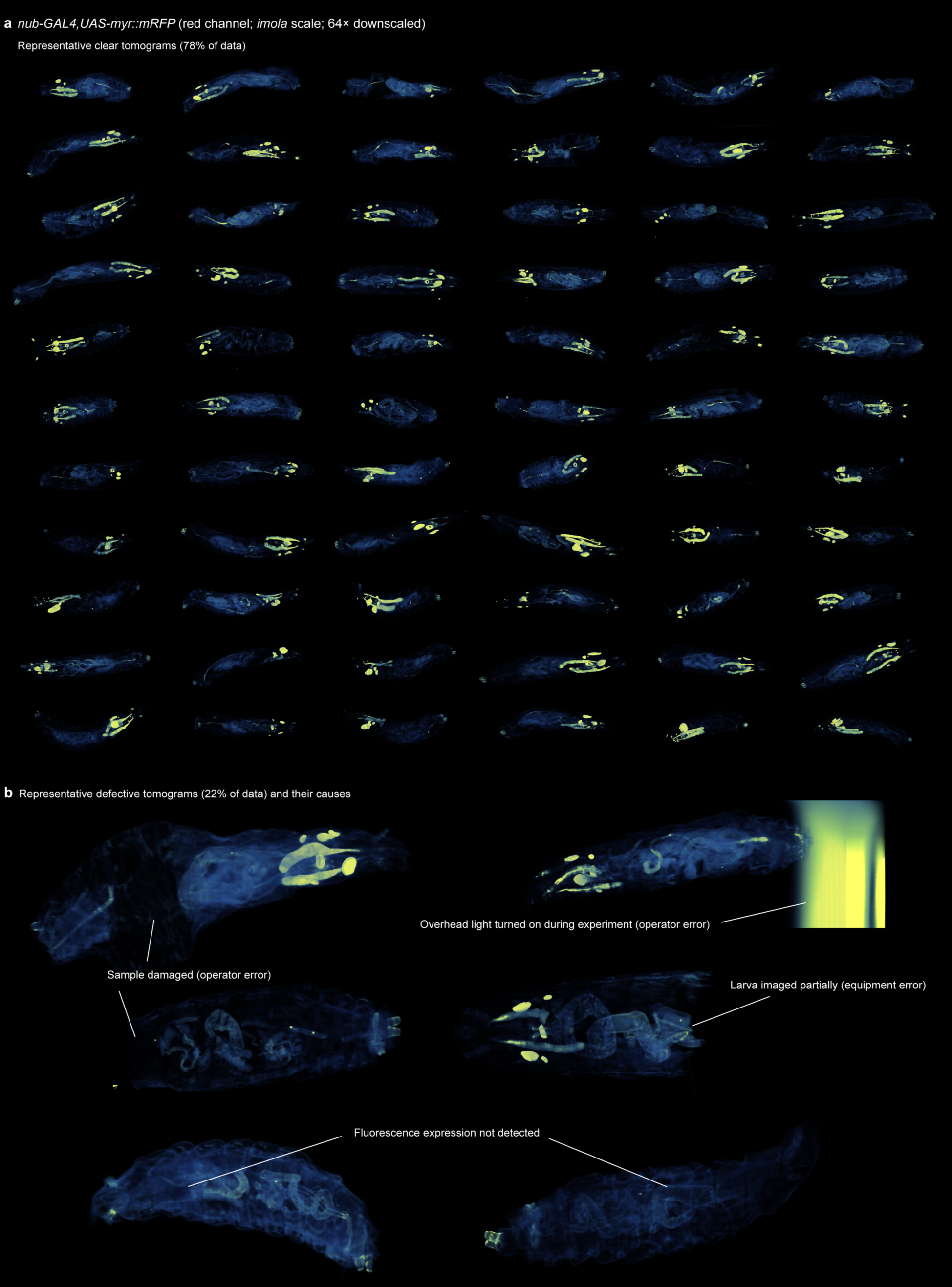
Library of 3D tomograms including causes of defective data. Shown are representative data obtained from flow zoometry of *nub-GAL4* larvae in a single experiment (*n* = 123). **a,** Representative 3D tomograms of *nub-GAL4* larvae which we designated as “excellent” (*n* = 96). In all clear tomograms, all 7 target tissues can be clearly identified against the autofluorescence background, and there are no noticeable defects. All tomograms in **a** are displayed as *xz* orthographic projections at the same intensity, color (*imola*), and size scales. Evident is variation in the autofluorescence levels, developmental stages, and positions of the larvae. **b,** Representative defective 3D tomograms (*n* = 27) with several types of defects. The defects shown are intended to represent commonly-encountered issues in flow zoometry. These include operator errors such as damage to samples and improper handling of equipment (e.g., turning on overhead lights), equipment errors such as partial data acquisition (i.e., the flow zoometer did not scan the entire larva), and problems of unknown causes such as a lack of fluorescence expression in samples. All tomograms in **b** are displayed as *xz* orthographic projections at the same intensity, color (*imola*), and size scales.

**Supplementary Figure 10.**
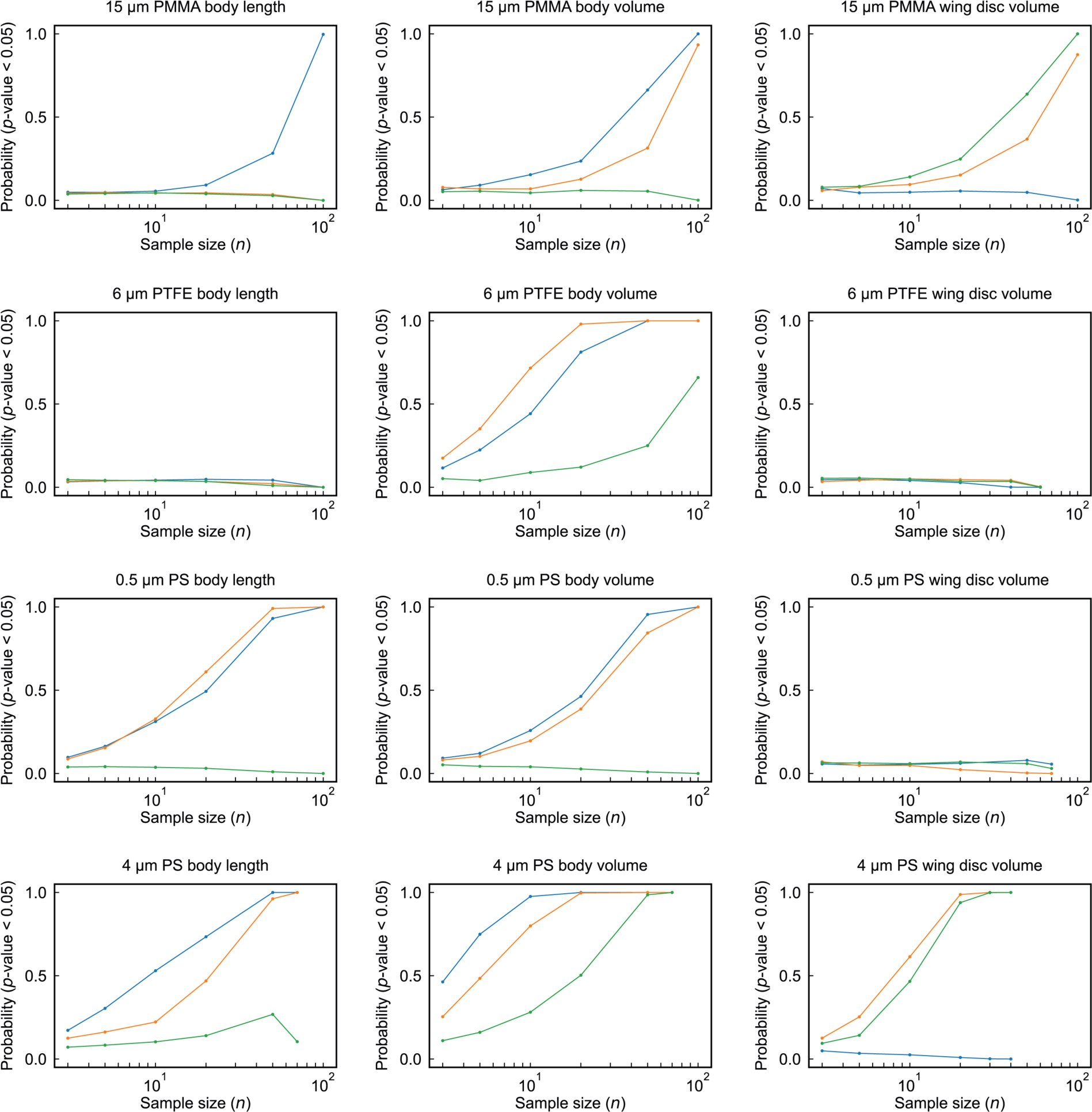
Monte Carlo simulations of statistical significance probability in toxicology screening. Each plot depicts the results of Monte Carlo simulations of *p*-value calculations based on random sub-sampling of the data from the toxicology screening experiments. As the sample size (*n*) for each *p*-value calculation is increased, the probability of detecting a significant result (defined here as *p* < 0.05) increases, supporting the importance of flow zoometry as a tool for large-scale biological assays. For each data point, 1,000 simulations were performed.

