## Supplementary Information for "Flow zoometry of *Drosophila*"

**Supplementary Table 1 | Throughput of large-scale whole-animal toxicological screening**

| Sample | Raw images | *n_1_* for body length | *n_2_* for body volume | *n_3_* for wing discs | Clearing time (hr) | Loading, flowing, and imaging time (min) | Loading, flowing, and imaging throughput (fly/min) | Total cohort time (hr) |
| --- | --- | --- | --- | --- | --- | --- | --- | --- |
| 15 µm PMMA control | 150 | 149 | 141 | 190 | 21 | 163 | 0.91 | 28 |
| 15 µm PMMA 0.25 µg/g | 134 | 129 | 114 | 162 |  | 140 | 0.93 |  |
| 15 µm PMMA  2.5 µg/g | 119 | 119 | 109 | 156 |  | 120 | 0.98 |  |
| 6 µm PTFE control | 109 | 108 | 105 | 102 | 21 | 136 | 0.80 | 28 |
| 6 µm PTFE 0.25 µg/g | 106 | 106 | 104 | 125 |  | 118 | 0.90 |  |
| 6 µm PTFE  2.5 µg/g | 120 | 120 | 114 | 116 |  | 168 | 0.71 |  |
| 0.5 µm PS control | 110 | 110 | 110 | 156 | 21 | 114 | 0.96 | 28 |
| 0.5 µm PS 0.26 µg/g | 112 | 112 | 109 | 148 |  | 136 | 0.82 |  |
| 0.5 µm PS 2.6 µg/g | 122 | 121 | 117 | 139 |  | 173 | 0.71 |  |
| 4 µm PS control | 77 | 73 | 71 | 114 | 21 | 77 | 1.0 | 26 |
| 4 µm PS 0.26 µg/g | 109 | 107 | 102 | 46 |  | 138 | 0.79 |  |
| 4 µm PS 2.6 µg/g | 84 | 83 | 80 | 66 |  | 96 | 0.88 |  |

**Supplementary Table 2 | Statistical results of large-scale whole-animal toxicological screening**

| Sample | Body length (mm) | Standard deviation (mm) | Body volume (mm^3^) | Standard deviation (mm^3^) | Total wing disc volume (mm^3^) | Standard deviation  (mm^3^) |
| --- | --- | --- | --- | --- | --- | --- |
| 15 µm PMMA control | 4.030 | 0.510 | 0.833 | 0.346 | 0.880 × 10^-3^ | 0.549 × 10^-3^ |
| 15 µm PMMA 0.25 µg/g | 3.883 | 0.431 | 0.691 | 0.303 | 0.953 × 10^-3^ | 0.625 × 10^-3^ |
| 15 µm PMMA  2.5 µg/g | 3.841 | 0.465 | 0.709 | 0.285 | 0.710 × 10^-3^ | 0.478 × 10^-3^ |
| 6 µm PTFE control | 3.790 | 0.548 | 0.898 | 0.196 | 1.019 × 10^-3^ | 0.732 × 10^-3^ |
| 6 µm PTFE 0.25 µg/g | 3.702 | 0.489 | 0.715 | 0.212 | 1.016 × 10^-3^ | 0.672 × 10^-3^ |
| 6 µm PTFE  2.5 µg/g | 3.640 | 0.681 | 0.655 | 0.229 | 0.884 × 10^-3^ | 0.696 × 10^-3^ |
| 0.5 µm PS control | 3.685 | 0.462 | 0.712 | 0.178 | 0.882 × 10^-3^ | 0.512 × 10^-3^ |
| 0.5 µm PS 0.26 µg/g | 3.372 | 0.540 | 0.592 | 0.208 | 0.807 × 10^-3^ | 0.511 × 10^-3^ |
| 0.5 µm PS 2.6 µg/g | 3.332 | 0.453 | 0.596 | 0.249 | 0.886 × 10^-3^ | 0.553 × 10^-3^ |
| 4 µm PS control | 3.353 | 0.360 | 0.802 | 0.199 | 0.759 × 10^-3^ | 0.459 × 10^-3^ |
| 4 µm PS 0.26 µg/g | 3.100 | 0.466 | 0.496 | 0.175 | 0.691 × 10^-3^ | 0.498 × 10^-3^ |
| 4 µm PS 2.6 µg/g | 3.062 | 0.410 | 0.560 | 0.168 | 0.411 × 10^-3^ | 0.274 × 10^-3^ |

**Supplementary Table 3 | Tissue-clearing reagent administration in full protocol**

| Purpose | Reagent | Exposure time | Note |
| --- | --- | --- | --- |
| Dechorionation | 4% bleach | 10 min | Manual, optional |
| Washing | Distilled water | 5 min | Automated |
| Washing | Distilled water | 5 min | Automated |
| Washing | Distilled water | 5 min | Automated |
| Fixation | 4% PFA | 120 min  (room temperature) | Automated |
| Delipidation/washing | 0.2% PBT | 90 min | Automated |
| Delipidation/washing | 0.2% PBT | 90 min | Automated |
| Delipidation/washing | 0.2% PBT | 90 min | Automated |
| Dehydration | 10% 1-propanol | 120 min | Automated |
| Dehydration | 30% 1-propanol | 120 min | Automated |
| Dehydration | 50% 1-propanol | 120 min | Automated |
| Dehydration | 70% 1-propanol | 120 min | Automated |
| Dehydration | 100% 1-propanol | 60 min | Automated |
| Dehydration | 100% 1-propanol | 60 min | Automated |
| RI matching | Ethyl cinnamate | 240 min | Automated |

**Supplementary Table 4 | Timing of tissue-clearing reagent administration in split protocol**

| Purpose | Reagent | Exposure time | Note |
| --- | --- | --- | --- |
| Dechorionation | 4% bleach | 10 min | Manual, optional |
| Washing | Distilled water | 5 min | Manual |
| Washing | Distilled water | 5 min | Manual |
| Washing | Distilled water | 5 min | Manual |
| Fixation | 4% PFA | Overnight (4 °C) | Manual |
| Delipidation/washing | 0.2% PBT | 90 min | Automated |
| Delipidation/washing | 0.2% PBT | 90 min | Automated |
| Delipidation/washing | 0.2% PBT | 90 min | Automated |
| Dehydration | 10% 1-propanol | 120 min | Automated |
| Dehydration | 30% 1-propanol | 120 min | Automated |
| Dehydration | 50% 1-propanol | 120 min | Automated |
| Dehydration | 70% 1-propanol | 120 min | Automated |
| Dehydration | 100% 1-propanol | 60 min | Automated |
| Dehydration | 100% 1-propanol | 60 min | Automated |
| RI matching | Ethyl cinnamate | 240 min | Automated |

**Supplementary Method 1 | Details on 3D U-Net image segmentation**

Our algorithm introduces a robust preprocessing pipeline tailored to the extraction of discrete larval structures from 3D tomograms obtained by flow zoometry. After multi-class segmentation, quantitative analysis of each tissue class can be applied to biological applications. Our pipeline consists of the following sequential steps:

(1) Noise reduction and background elimination. In the initial stage, we implemented an adaptive thresholding technique (Otsu’s method) to differentiate between the larva and the background within an intensity range of 100-1000. This was complemented by Gaussian blurring to smooth the image, minimizing the effects of noise and artifacts. After thresholding, we employed a connectivity analysis to identify and retain the largest connected component, presumed to be the larva body, effectively isolating it from the surrounding noise.

(2) Model training with nnU-Net pipeline. For segmentation, we used a standard nnUNet pipeline, opting for a 3D full-resolution model. The model was strengthened via a 5-fold cross-validation (with all annotations used) to ensure generalizability. An ensemble of the trained models was then used to enhance the predictive performance on the test set, followed by a tailored post-processing step to refine the segmentation results.

(3) Feature analysis for larval characterization. The segmented larval structures underwent a feature analysis to extract biological insights. Notably, the larval length was ascertained by first computing the skeletal representation of the larva and then calculating the maximum Feret diameter of the skeleton. This approach significantly reduced computational demands while providing accurate morphometric assessments.

**Supplementary Method 2 | Details on image preprocessing, wing disk region segmentation, feature extraction, and classification in the Random Forest classifier**

The following preprocessing pipeline was designed to transform the distribution of intensity of the *ptc* regions of wing disc tissues into binary segmentations.

(1) Bounding box estimation. The goal of bounding box estimation is to accurately define the spatial extent of the larva within the 3D dataset, isolating it from the surrounding capillary structure. An orthogonal coordinate system was defined so that the long axis of the capillary was the *z* axis while its two orthogonal directions were the *x* and *y* axes. For each 2D *xy* slice at each *z* the 3D dataset, the skewness $S_{x-y}$ of the fluorescence intensity distribution was estimated, resulting in the function $S_{x-y}(z$). This function was then binarized into $B(z)$using a threshold value of 0.5 according to the following rule:

$$B(z)= \left\{ \begin{aligned} 1\mathrm{if} S_{x-y}\left( z \right)>0.5 \\ 0\mathrm{if} S_{x-y}\left( z \right)\leq0.5 \end{aligned} \right.$$

The skewness can identify the region of larvae along *z* because 2D slices without larval presence were composed of comparatively weak fluorescence intensities which were close to symmetric, resulting in small skewness, while 2D slices with larval presence had high fluorescence intensity regions distinct from the weak background, resulting in a positively-skewed fluorescence intensity distribution across the whole 2D slice. Also, the skewness is free from influence from the absolute value of fluorescence intensity and is thus more robust than absolute intensity thresholding methods, as baselines may change across experiments or when implementing different fluorescence markers. Our skewness threshold of 0.5 was chosen based on its high success rate in discriminating larvae from background.

To determine the *z* slices where the larva body is likely to be present, an iterative algorithm was used to detect the longest contiguous sequence of 1 values in $B\left( z \right)$. This skewness estimation followed by the binarization and the detection of the longest sequence of 1 values was similarly applied in the *x* and *y* directions. By identifying the coordinates of the longest sequences of 1 values in the $x$, $y,$ and $z$directions, we estimated a 3D bounding box around the body of the larva.

(2) Larva body segmentation. Following estimation of the bounding box, the data within the bounding box underwent a 3D Gaussian blurring filter with a standard deviation of 1.5. The choice of 1.5 corresponds to a kernel size of dimensions $(x,y,z)=7\times7\times7$ so that the intensity value of each voxel was replaced by a gaussian weighted average taken over the nearest 343 voxels, to eliminate high-frequency noise but leave the main structural features of larvae intact. Subsequently, a Gaussian mixture model (GMM) fitted with 3 components was applied to the denoised voxels to segment the larva body within the bounding box. Within the bounding box, the cluster of voxels exhibiting the lowest average intensity value was assigned as background where the larva did not exist and was labeled as 0. The remaining two clusters characterized by higher average intensities were together recognized as the larva body region and labelled as 1.

(3) Cropping of the larva head containing salivary glands and wing discs. Following the larva body segmentation, the 99th percentile of voxel intensities is estimated within the larva body region, $Q_{(I_{99}\%)}$. This value was utilized as a threshold to detect the salivary glands and wing discs, which are expected to be the brightest regions of the flies. We then excluded all isolated binary shapes smaller than 3,500 voxels (the average body size of larvae was 2,102,802 voxels in this study, such that 3,500 voxels corresponded to 0.16 % of the larva body, on average). Here we identified the highest-intensity regions as salivary glands which must be excluded in later analysis. We found that the removal of isolated binary shapes smaller than 3,500 voxels left the largest segments corresponding to salivary glands.

Salivary glands and wing discs are anatomically close to each other, such that the salivary gland positions and wing disc positions along the *z* axis were similar. To pinpoint the region of interest along the *z* axis, a function $L(z)$ was applied to each *z*-slice. $L(z)$ represents the number of pixels belonging to the isolated, bright segments that are anticipated to include salivary glands and wing discs.$L(z)$ was then transformed into a binary representation: 1 if $L\left( z \right)>0$*,* and 0 otherwise. Since the salivary glands and wing discs are anatomically closer to the head than the tail, the orientation of a larva along the *z* axis was estimated by the function $L(z)$ by comparing its value over the intervals of $[1, N_{z}/2]$ and $[N_{z}/2+1, N_{z}]$, where $N_{z}$ represents the total number of *z*-slices in the bounding box. A recursive algorithm then identifies the longest sequence of 1 values in $L(z)$, defining the interval of interest as $[z_{\min}, z_{\max}]$. Depending on the larva’s head orientation, the region of interest which includes the salivary glands and wing discs tissues is defined as follows: [$z_{\min}-40, z_{\max}+30]$ if the head is toward positive *z*; [$z_{\min}-30, z_{\max}+40]$ if the head is toward negative *z* . Our choices for the segmented range were empirically determined, such that wing discs were segmented in their entirety. This is our primary definition of the tentative wing disc, “wing disc”, region to be refined and segmented further.

(4) Segmentation of the wing discs. After identifying the head region along the *z* axis, the cropped 3D dataset of the estimated larval head region was initially denoised by an Oracle-based 3D Discrete Cosine Transform (OCDT3D) filter with parameters $\left( \sigma,\tau\right)=(15,15)$^1^. The voxel intensities $I(x,y,z)$ of the region were transformed through a series of steps to enhance the comparability across different measurement dates and different batches. The voxels were initially subject to a logarithmic transformation to reduce the dynamic range of the voxel intensities, to mitigate the impact of outliers. This was followed by min-max normalization and a global contrast normalization. Next we identified and isolated the top 25% brightest voxels to remove major autofluorescence noise. The remaining voxels were clustered into 2 groups using the *k-*means algorithm, based on Euclidean distance. The cluster with the highest average intensity provided a preliminary estimation of the segmented tissues, including the wing disc tissues. To refine the segmentation, postprocessing steps were implemented. The first step filtered out segmented tissues that overlapped more than 12% (empirically chosen) with the salivary gland positions determined earlier. The previously estimated salivary gland voxels were required to be contiguous over 3,500 voxels, but the present clustering process was based on the cropped localized regions and resulted in refined, larger segments corresponding to salivary glands in addition to wing disc segment. This process was followed by the generation of a 2D projection of each remaining segmented tissue on the *xy* plane, followed by the discarding of any segments with a projection circularity less than 0.35 (corresponding to debris). Subsequently, all isolated shapes with a longest principal axis of less than 20 voxels were removed. The final stage of the segmentation is dependent on the number of isolated tissues remaining. If more than three isolated shapes remained, a *k-*means clustering with $k=3$ was performed, using the centroid coordinates and average voxel intensity values of each remaining shape as features. The cluster displaying the highest average intensity was then assigned as the final segmentation of the wing discs. However, if there were three or fewer isolated shapes remaining, these sets of tissues were considered as the final segmentation without further clustering.

**Feature extraction and Random Forest classifier**

From the binary segmentation of the wing discs, we obtained a set of shapes labelled by a connected component algorithm. Next, we described this set of connected components by computing both the average values and the maximum values of distinct features: volume, bounding box dimensions (width, height, depth), solidity (volume to convex hull volume ratio), principal axis length ratios (1^st^/2^nd^ and 1^st^/3^rd^) of the approximating ellipsoid, compactness (volume to surface area ratio), and the ratio of bounding volume to the component's volume. Additionally, we employed the D2 shape descriptor, which is based on the distribution of distances among voxels within each shape. We quantified this distribution of distance by estimating the quantiles 0.05,0.25,0.5,0.75,0.95 and 1. The segmented larva wing discs were described by a set of 30 features which were then used as inputs for a Random Forest algorithm to classify the wing discs segmentation between ptc*>mCherry 4-hit and* *ptc>mCherry.* Importantly, our extracted morphometric features of wing discs from 3D fluorescent images were coordinate-free, such that data were independent of the orientation of larva and any data augmentation was not required. The Random Forest classifier was tuned and trained by using a nested cross validation approach with data organized by 6 measurement dates. The performance of the model’s generalization was evaluated using several metrics such as the area under the receiver operating characteristic curve ($AUC$), $\mathrm{accuracy}= \frac{TP+TN}{TP+TN+FP+FN}$ , $\mathrm{recall}= \frac{TP}{TP+FN}$, $\mathrm{precision}=\frac{TP}{TP+FP}$ , and $F1-score= \frac{2\times precision\times recall}{precision+recall}$ , where $TP$ is true positives, $TN$ is true negatives, $FP$ is false positives, and $FN$ is false negatives.

**Supplementary Discussion** **1** **| Additional possible directions for developing the flow zoometer**

Infrequent motion blur during imaging could be mitigated by installing custom-made pinch valves to better ensure the stoppage of flow, or by fine-tuning the viscosity and density of the flowing liquid to better immobilize the sample. Another example is the system of picking individual larvae in the dispersing chamber, which could be changed from a stochastic process to a more directed process by installing a more sophisticated jet mechanism and pick-up method for high-precision flow control and larva selection. As a final example of a potential technical improvement, by changing the LSM from single-sided to two-sided illumination, it should be possible to minimize fluorescence intensity gradient effects in the sample while using ultraviolet and blue excitation sources.

**Supplementary Note 1** **| Fluorescence patterns of 4-color larvae**

In Figure 2a we show 4-color tomograms of *UAS-mtdTomato,10xQUAS-6xGFP/nub-GAL4;r4-QF2^3xP3-RFP^/+* with DAPI staining. It should be noted that the fluorescent protein expression patterns in the orange and red channels are expected to be highly similar due to the spectral overlap between mtdTomato and RFP. Our results shown in the figure are consistent with this expectation; the displayed orange and red 3D tomograms share similar features. Much of their difference can be attributed to the difference in the level of autofluorescence background, which is stronger in the orange channel as compared to the red channel.
